# Molecular Mechanism of Active Cas7-11 in Processing CRISPR RNA and Interfering Target RNA

**DOI:** 10.1101/2022.06.22.497266

**Authors:** Hemant Gowswami, Jay Rai, Anuska Das, Hong Li

**Affiliations:** Institute of Molecular Biophysics, Florida State University, Florida State University, Tallahassee, FL, USA; Department of Chemistry and Biochemistry, Florida State University, Tallahassee, FL 32306, USA

## Abstract

Cas7-11 is a Type III-E CRISPR Cas effector that confers programmable RNA cleavage and has potential applications in RNA interference. Cas7-11 encodes a single polypeptide containing four Cas7- and one Cas11-like segments that obscures the distinction between the multi-subunit Class 1 and the single-subunit Class-2 CRISPR-Cas systems. We report a cryo-EM structure of the active Cas7-11 from *Desulfonema ishimotonii* (DiCas7-11) that reveals the molecular basis for RNA processing and interference activities. DiCas7-11 arranges its Cas7- and Cas11-like domains in an extended form that resembles the backbone made up by four Cas7 and one Cas11 subunits in the multi-subunit enzymes. Unlike the multi-subunit enzymes, however, the backbone of DiCas7-11 contains evolutionarily different Cas7 and Cas11 domains, giving rise to their unique functionality. The first Cas7-like domain nearly engulfs the last 15 direct repeat nucleotides and is responsible for processing and recognition of the CRISPR RNA. Whereas both the second and the third Cas7-like domains mediate target RNA cleavage, they differ in metal requirement for catalysis. The long variable insertion to the fourth Cas7-like domain has little impact to RNA processing or targeting, suggesting the possibility for engineering a compact and programmable RNA interference tool.

**One Sentence Summary:** Structures of Cas7-11 reveal the molecular basis for processing CRISPR RNA and for cleaving target RNA.

## Main Text

The CRISPR-Cas systems confer adaptive immunity to prokaryotic hosts against invading viruses by encoding a range of different CRISPR-Cas efforts that interfere with the invader nucleic acids ^1^. Three types of CRISPR-Cas effectors are known to utilize programmable CRISPR RNA (crRNA) to guide cleavage of the complementary target RNA. The multi-subunit Type III effectors, exemplified by the Type III-A (Csm) and the III-B subtypes (Cmr), assemble 4-5 Cas7, 2-3 Cas11, and 1 Cas10 subunits into a sea horse-shaped helical enzyme ^2^. They cleave the target RNA at a 6-nucleotide (nt) interval within the complementary region that coincides with the evenly spaced Cas7 subunits ^3–5^. The Type VI, or Cas13, is a single subunit and substantially smaller effector. Unlike Csm/Cmr, Cas13 employs two Higher Prokaryotic and Eukaryotic binding domains (HEPN) in cleaving the target RNA outside the complementary region ^6, 7^. The recently discovered Type III-E effector, Cas7-11 or gRAMP (for giant Repeat Associated Mysterious Protein), is also a single subunit effector with fused Cas7 and Cas11 segments ^8, 9^ (Figure 1a). Unlike Cas13 but similar to Csm/Cmr, Cas7-11 employs the Cas7-like segments to cleave crRNA-guided target RNA (Figure 1a). Interestingly, whereas Cas13 can distinguish self from foreign RNA by utilizing the 3’ protospacer flanking sequence (PFS) in the target RNA ^10^, both Csm/Cmr and Cas7-11 are insensitive to 3’ PFS in cleaving their respective target RNA ^8, 9^. The three effectors also differ in crRNA processing. Csm/Cmr utilize an independent processing endonuclease, Cas6, to result in a mature crRNA containing an 8-nucleotide (nt) repeat (5’-tag) linked to the spacer ^11^. By contrast, both Cas13 and Cas7-11 process their own crRNA ^7–9^. Cas7-11 is therefore believed to be an evolutionary intermediate between the Class 1 and 2 effectors. Interestingly, Cas7-11 has been demonstrated form a complex with the caspase-like TPR-CHAT peptidase, suggesting a potential for a viral RNA-induced and protease-mediated antiviral immunity ^8^. Given the known collateral nuclease activities of Cas13 ^6, 7^ and Csm/Cmr ^3, 4, 12^, and the complex enzyme composition of Csm/Cmr, Cas7-11 provides a desirable platform to further develop RNA interference and editing tools. To understand the molecular basis for crRNA processing and target interference of Cas7-11, we determined a cryo-electron microscopy (cryo-EM) structure of *Desulfonema ishimotonii* Cas7-11 (DiCas7-11) at an overall resolution of 2.82 Å (Figure 1b-1c, Supplementary Figure 1, Supplementary Figure 1 & Supplementary Table 1). DiCas7-11 has been demonstrated to function in programmable RNA cleavage and editing both in vitro and in mammalian cells ^8, 9^. Our structure provides the architecture of the enzyme and the molecular basis for its enzymatic activities.

**Figure 1.**
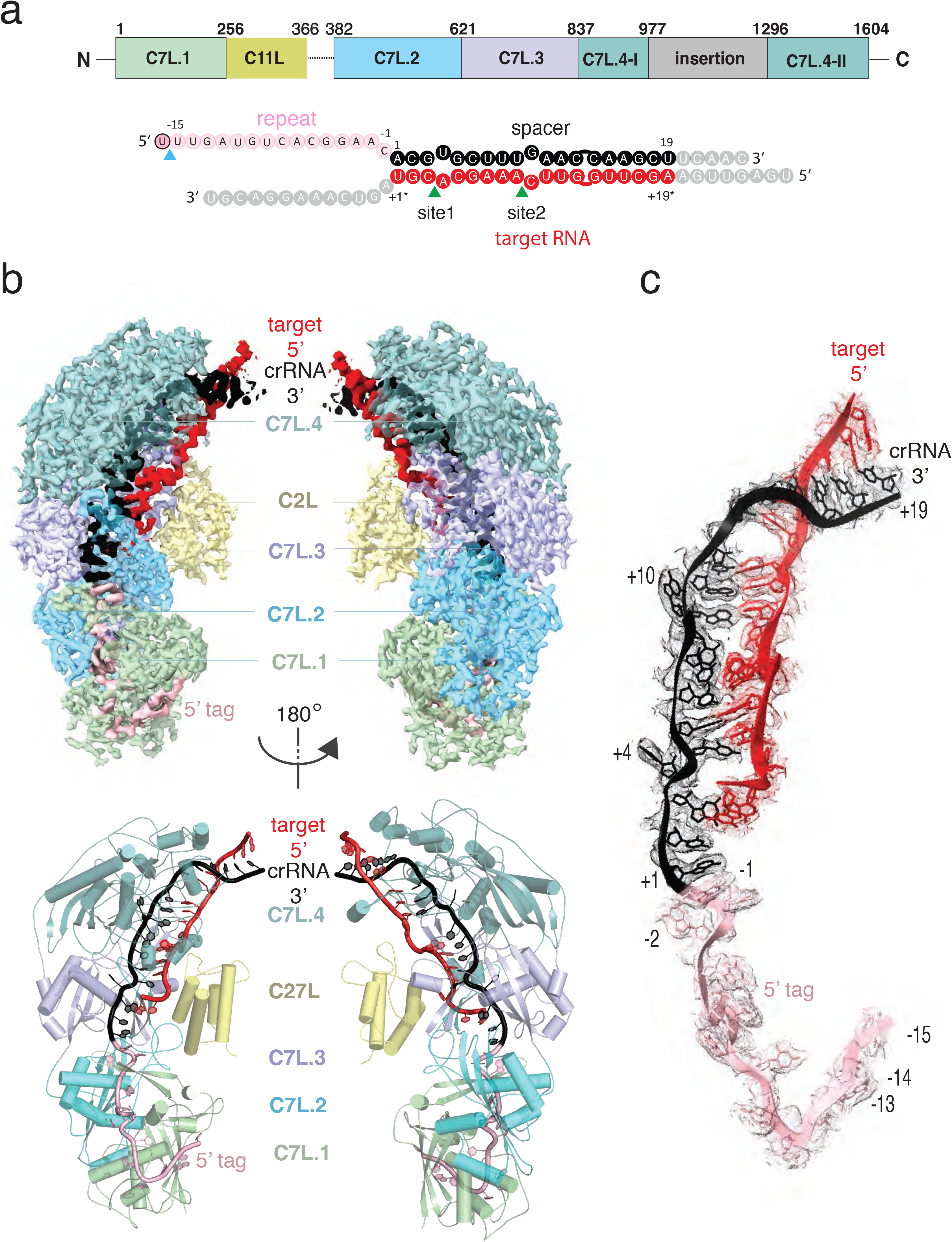
Structure overview of DiCas7-11-crRNA-target-RNA ternary complex. (a) Domain organization of DiCas7-11 and schematic representation of crRNA-target RNA duplexes used in the study. The blue and green colored triangles indicate the pre-crRNA processing and target RNA cleavage sites, respectively. C7L denotes Cas7-like domain and C11L denotes Cas11-like domain. (b) Top, electron potential density map of DiCas7-11-crRNA-target-RNA ternary complex shown in two different orientations. Bottom, cartoon representation of DiCas7-11-crRNA-target-RNA ternary complex shown the same views as in top panel with corresponding colors representing protein domains and the two RNA strands. (c) Close-up view of the density for the crRNA (spacer:black, repeat:light pink) and target RNA (red) duplex.

The wild-type DiCas7-11 was incubated with its precursor crRNA and a complementary target RNA under a reactive condition before being made frozen specimen (Supplementary Figure 1). Under this condition, DiCas7-11 successfully processes the precursor crRNA and cleaves the target RNA (Supplementary Figure 1). The density map resolves most of the DiCas7-11 protein, the crRNA and the partially cleaved target RNA (Figure 1b-1c, Supplementary Figure 2 & Supplementary Figure 3). The core Cas7-11 assembly is half-moon shaped with four Cas7-like (C7L) domains (C7L.1 – C7L.4 from N- to the C-terminus) forming a long ridge and a single Cas11-like domain (C11L) occupying the crescent center. The entire DiCas7-11 complex can be superimposed onto the closely matched homologous *Lactococcus lactis* Csm (LlCsm) complex ^13^ with the C7L.1-C7L.4 ridge matching that formed by four Csm3 subunits and C11L matching the Csm2 subunit adjacent to Cas10 (Supplementary Figure 4). The similarity in the bound RNA trajectory at the core between the two complex is striking (Supplementary Figure 4), suggesting a similar RNA binding apparatus in both complexes. DiCas7-11 lacks the domains equivalent to the Csm4 and the Csm1 subunits and, thus, the functions associated with them. In Csm complexes, Csm4 recognizes and stabilizes the 8-nt 5’-tag derived from the direct repeat whereas Csm1 carries out cyclic oligoadenylate synthesis and ancillary DNA cleavage ^13–15^. Csm1 also secures the 3’-PFS of the cognate target RNA, and thus, plays a role in discrimination of self from foreign RNA^13–15^.

Despite the overall structural similarity between the C7L- and the LlCsm-formed ridge (Supplementary Figure 4 & Supplementary Figure 5), each C7L differs slightly in protein sequence and folding (Supplementary Figure 4b), which gives rise to their different roles in binding and cleaving RNA. The 34-nt crRNA lies along the C7L-ridge with its 5’ tag spanning C7L.1-C7L.2 and the spacer region covering C7L.3-C7L.4 (Figure 1b-1c & Supplementary Figure 3). The first 18 nucleotides of the target RNA (+19* ∼ +2*) remain base paired with the spacer region of the crRNA (Figure 1c, Supplementary Figure 3, & Supplementary Figure 5). The short span of the guide-target region on C7L domains explains the two, instead of four as in Csm, sites of target cleavage ^8, 9^.

The observed structure suggests that the precursor crRNA (pre-crRNA) is processed by the first C7L domain, which yields a mature crRNA containing the last 15 nucleotides of the direct repeat linked to the programmed spacer (Figure 1c, Figure 2a, & Supplementary Figure 3). To confirm the site of processing, we subjected synthetic pre-crRNA containing 2’-deoxy modification at -16, -15, or -14 position to the processing reaction, respectively, and found that the cleavage products are consistent with the density-derived 15-nt 5’-tag (Figure 2b & Supplementary Figure 3). Strikingly, the sequence identity downstream of U(−15) is not important for processing, as those substituted with poly-adenine or the previously characterized *Candidatus Scalindua broadae* (Csb) pre-crRNA are successfully processed by DiCas7-11 (Figure 2c). A well conserved histidine residue, His43, is immediately next to the leaving 5’-hydroxyl oxygen of the -15 nucleotide, suggesting its role in processing. Consistently, His43 to alanine mutation (H43A) abolished pre-crRNA cleavage (Figure 2c). The requirement for His43 with no divalent metals in processing suggests that the C7L.1 employs a RNase A-like mechanism similar to that suggested for Cas6 ^16, 17^. Interestingly, CsbCas7-11 contains threonine in place of histidine and does not process its crRNA at position -15 ^8^, suggesting that precise processing may not be required for the RNA interference activity.

**Figure 2.**
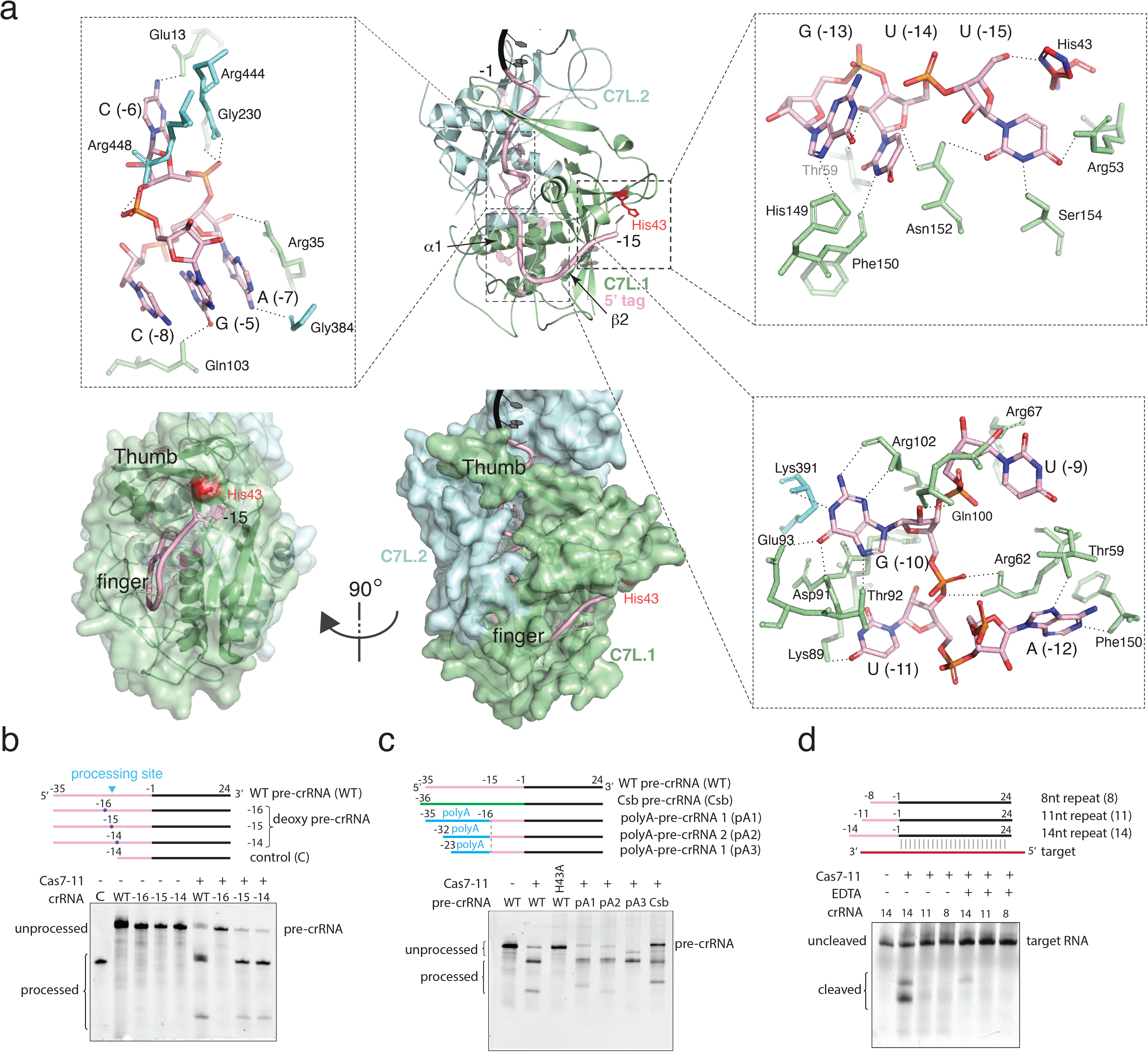
Precursor crRNA processing and recognition. (a) The mode of Cas7-like domain 1 (C7L.1) and C7L.2 interaction with the processed crRNA nucleotides -15 to -1 in both cartoon (Top) and surface (bottom) representations. Key secondary elements involved in crRNA interaction are labeled. Insets indicate close-up views around U(−15)-U(−13)-G(−13), the tight RNA turn, and the conserved A(−12)-U(−11)-G(−10)-U(−9) tetranucleotide. Dashed lines indicate close polar contacts. (b) RNA processing results analyzed on polyacrylamide urea gel with the wild-type and deoxy modified precursor crRNA (pre-crRNA) by DiCas7-11. The sites of deoxy modification are indicated by marked nucleotide positions on the wild-type pre-crRNA, respectively. The control RNA contains the last 14 nucleotides of the repeat plus the spacer. Pre-crRNA processing site is indicated by a blue triangle. (c) RNA processing results analyzed on polyacrylamide urea gel with the wild-type and the His43 to alanine mutant of DiCas7-11 with the wild-type, poly-adenine-substituted, and the *Candidatus Scalindua broadae* (Csb) pre-crRNA. (d) Target RNA cleavage results analyzed on polyacrylamide urea gel with the wild-type and truncated precursor crRNA (pre-crRNA) by DiCas7-11 in the presence and absence of EDTA.

The processed 15-nt 5’-tag interacts extensively with C7L.1 and to less extend with C7L.2 (Figure 2a & Supplementary Figure 5). The protein nearly buries the entire 5’-tag and thus precluded it from further base pairing with a complementary RNA. The C7L.1 forms a ferredoxin fold (π2τπ3τα2π1τα1π4τ) highly abundant in CRISPR-Cas components ^18, 19^. It uses α1, π2, a long π-hairpin connecting π2 to π3 (thumb) and a C7L.1-specific insertion loop (finger) to secure the significantly bent 5’-tag, resembling a rope-gripping right hand (Figure 2a & Supplementary Figure 5). The first nucleotide, U(−15), forms a hydrogen bond network with a number of residues including the well conserved His43 (Figure 2a & Supplementary Figure 5). Removal of U(−15) did not impact target RNA cleavage (Figure 2d), suggesting the possibility that the U(−15)-protein interactions may be important for processing rather than target interference. The most extensive interactions take place at the strictly conserved A(−12)-U(−11)-G(−10)-U(−9) tetranucleotide. Both sidechain as well as mainchain atoms of C7L.1 participate in “reading” the four RNA bases. Strikingly, all edges of G(−10), the Watson-Crick, the Hoogsteen, and the Sugar, are in close contacts with the C7L.1 residues (Figure 2a). Downstream of the AUGU tetranucleotide is a tight right-handed helical turn formed by C(−8)-A(−7)-C(−6)-G(−5), reminiscent the 3_10_ helix in proteins. The turn is stabilized by both base stacking as well as an unusual network of intra-strand polar contacts. The base of G(−5) interdigitates those of C(−8) and A(−7) with Arg35 on top, leaving C(−6) protruding into the interior of the protein (Figure 2a). Whereas phosphate backbone atoms in A-form RNA do not engage in intra-strand contacts, those within the turn mediate numerous interactions (Figure 2a). The strictly conserved G(−5) forms the most intra-strand interactions. Its N2 atom contacts the non-bridging oxygen of A(−7) while its non-bridging oxygen forms hydrogen bond with 2’-OH of C(−8). Finally, the 2’-OH of G(−5) forms bifurcated contacts with the N7 atom of the strictly conserved A(−7) and G(−4). The rest of the 5’-tag is clamped down by the thumb of C7L.1 and α1 of C7L.2 analogously as by two Cas7 subunits in Csm complexes ^13^. Consistent with the extensive 5’-tag-protein interactions, removal of either first 4 or 8 5’-tag nucleotides abolished RNA-guided target cleavage (Figure 2d).

The spacer region of the crRNA is captured by the rest of the C7L ridge and base paired with the target RNA (Figure 3a). Three of the four C7L domains contain the characteristic “thumb” that secure the crRNA at two evenly spaced (6-nt) kinks with bases flipped. The duplex bound by C7L.4 creates an extra spacing between the G(+13*)-C(+13) and G(+14*)-C(+14) pairs but no base flipping. Like the multi-subunit Type III efforts, the kinked crRNA-target RNA duplex create bended sugar-phosphate backbone at the locations that coincide with the sites of cleavage. Consistently, the target RNA containing 2’-deoxy modification at A(+4*) (site 1) and C (+10*) (site 2) prevented formation of any cleavage product (Figures 3b & 3c). Two acidic residues, Asp429 and Asp654, were previously shown to be critical to cleavage at site 1 and site 2, respectively ^9^. Satisfactorily, they are found near each corresponding scissile phosphate with the carboxylate oxygen 4.2-6.7 Å from the leaving 5’-oxygen (Figure 3a). At both sites, the phosphodiester bond breakage is further assisted by the near “in-line” geometry of the nucleophilic 2’-oxygen, the scissile phosphate and the leaving 5’-oxygen (Figure 3a). Two residues from the C11L domain, Arg283 (site 1) and His306 (site 2), where Arg283 is better conserved than His306, are observed to stabilize the attacking nucleotides by stacking on their bases (Figure 3a).

**Figure 3.**
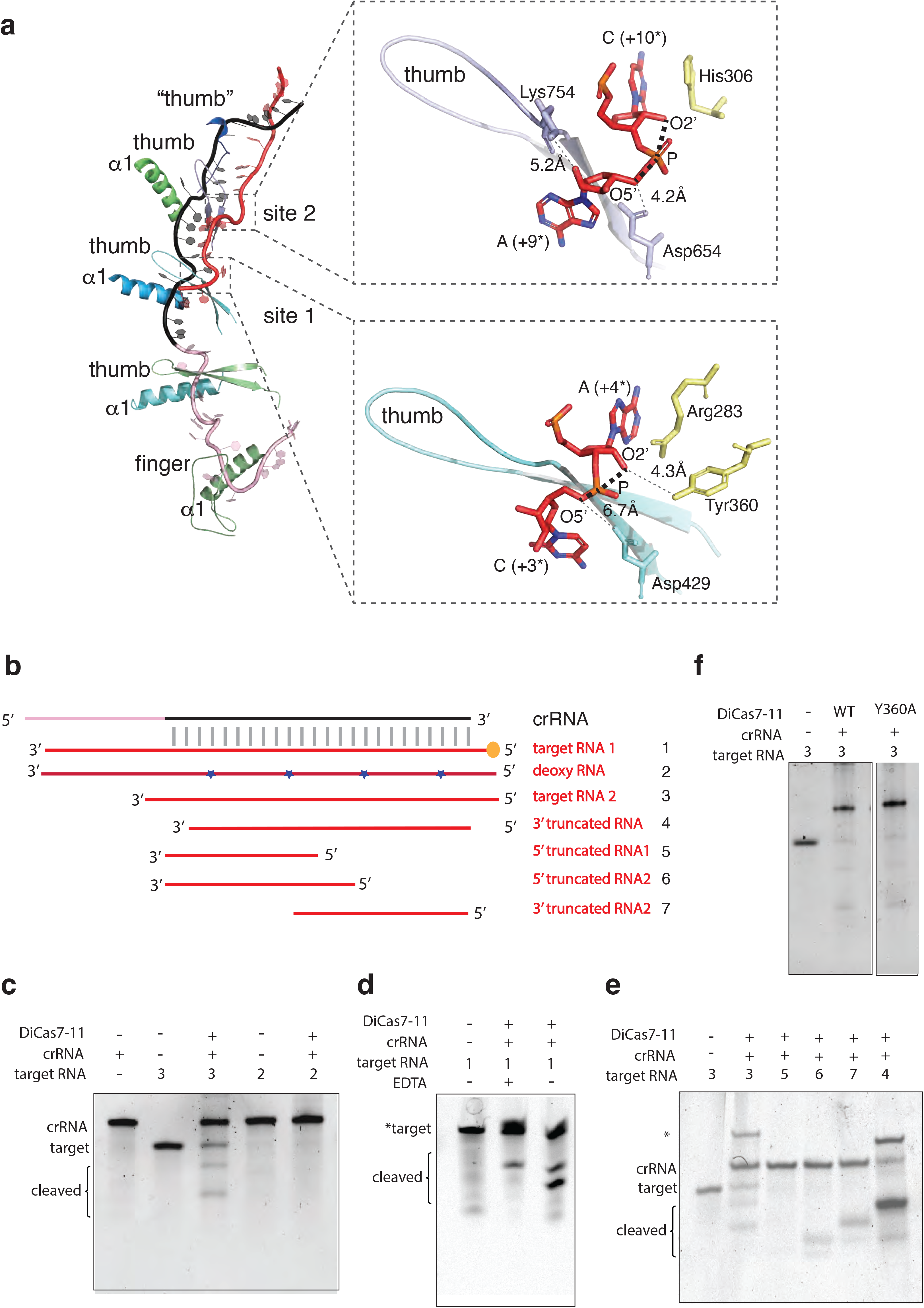
Target RNA cleavage mechanism. (a) Recognition of target RNA by the crRNA and DiCas7-11. The ferredoxin fold a1 and the thumb hairpin for each of the four Cas7-like (C7L) domains are shown as cartoons and colored as in Figure 1. “thumb” indicates the degenerate thumb feature for C7L.4 domain. The sites of target cleavage are boxed and shown in close-up views, respectively. Residues and amino acids are shown in stick models. The three atoms involved in the phosphodiester bond breakage are labeled and the angles they form are indicated by think dash lines. The closest of the three atoms to the putative catalytic residues, Asp654 (for site 2) and Asp429 (for site), are indicated by a connecting dash line. (b) Schematic of the labelled (*target) and non-labelled target RNA used to test DiCas7-11 cleavage activity. Asterisks mark the location of the deoxy modification on the target RNA (deoxy RNA). (c-f) Target RNA cleavage by DiCas7-11 and its Tyr360 to alanine mutant are analyzed on polyacrylamide urea gel. Location of the target RNA, crRNA, and cleaved target RNA products are marked. Asterisk in (f) indicates a possible nondenatured crRNA:target pair.

Interestingly in the homologous CsbCas7-11, Asp429 is not conserved and mutation of its equivalent Asp448 and other surrounding residues did not impact site 1 cleavage ^8^. We observed that, unlike cleavage at site 2 that strictly depends on Mg^2+^, cleavage at site 1 is independent of metal ions (Figures 2d, 3b & 3d), similar to its own processing activity or that by crRNA processing endonuclease Cas6 ^16, 17^. In Cas6, multiple amino acids are observed to fulfil the catalytic roles and therefore, the RNA geometry is believed to play a critical role in catalysis. It is thus possible that a correct RNA geometry at site 1, shaped by the protein, significantly accelerates catalysis. To access possible roles of other residues near site 1 in catalysis, we mutated the well conserved Tyr360 of C2L given its proximity to A23 (Figure 3a). Surprisingly, we found that Tyr360 is not required for site 1 cleavage (Figure 3b & 3f), indicating that, at least for DiCas7-11, Asp429 is sufficient for mediating the metal-dependent cleavage.

To access the length of base pairing required for target cleavage, we further examined the cleavage of a series of truncated target from either the 3’ or the 5’ end. We found that as short of 16 base pair in total length and 2 base pairs on one flanking end can facilitate RNA cleavage (Figures 3b & 3e), suggesting that protein and crRNA plays a significant role in shaping the target RNA for cleavage.

Though the final atomic model of DiCas7-11 lacks the large insertion to C7L.4 (residues 979-1297) due to weak density, focused classification and refinement led to a low-resolution map that matches the AlphaFold-predicted model of the insertion domain (Figure 4a & 4b). This model indicates that a large majority of the insertion domain is not engaged with any of the features described above and suggests the possibility that it is not essential to RNA-guided target cleavage. To test this hypothesis, we removed residues 1009-1220 to create DiCas7-11-Δint1. Consistently, we showed that DiCas7-11-Δint1 retains almost all RNA-guided target cleavage in an in vitro assay (Figure 4c).

**Figure 4.**
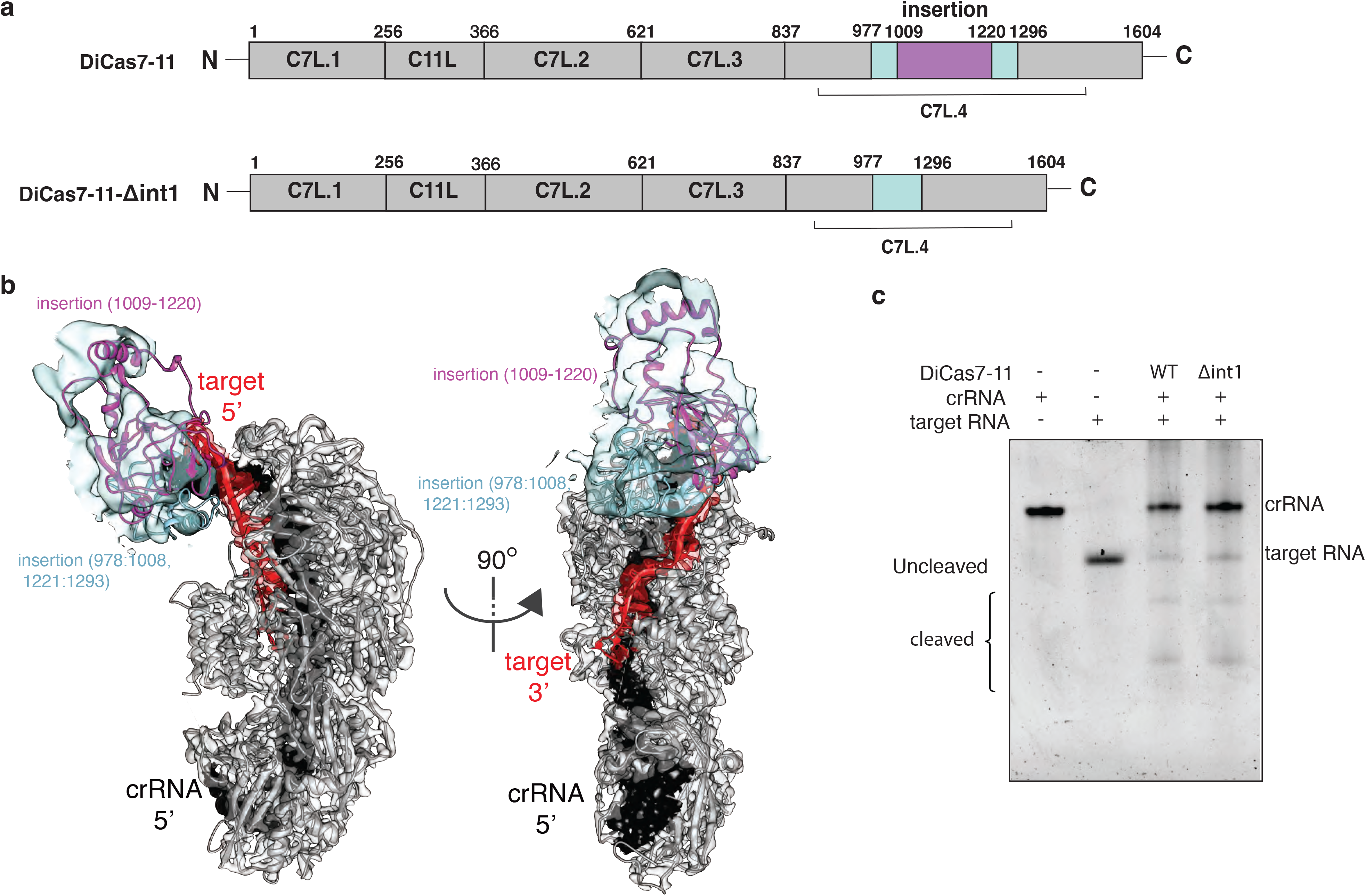
Engineering a compact DiCas7-11. (a) Schematic of domain organization of wild-type and an insertion deletion variant DiCas7-11-Δint1. The region removed is colored in purple and numbered. (b) Cartoon representation of DiCas7-11 overlaying with density map resulted from focused classification using a mask around the insertion domain. The insertion structure model is from AlphaFold prediction. The region removed is colored in purple and numbered. (c) Target RNA cleavage by DiCas7-11 and DiCas7-11-Δint1 are analyzed on a polyacrylamide urea gel. The crRNA, target and the cleaved products are labeled.

The structure and complementary biochemical assays show that Cas7-11 has a minimal architecture required for programmable RNA cleavage. The covalent linkage of the homologous units suggests an evolutionary advantage in dedicating Cas7-11 to RNA cleavage. Considering the known collateral RNase activity of Cas13 and the complicated Csm/Cmr systems, Cas7-11 offers a desirable alternative in developing gene regulation tools. While DiCas7-11 has been successfully demonstrated to function in mammalian cells, the efficiency and accuracy remain to be improved. With now the available structure and the accurately mapped processing and target cleavage sites, protein engineering may assist the efforts in designing improved Cas7-11-derived RNA interference platforms. DiCas7-1-Δint1 provides a proof-of-concept for such an effort.

## Methods and Materials

### Protein expression and purification

*Escherichia coli* NiCo21 (DE3) competent cells were transformed with the plasmid encoding sumo-tag fused DiCas7-11 (Addgene: 172503). A single colony was picked and transferred to 100mL LB media containing 50 μg/mL ampicillin and grown for 12 hours at 37°C before inoculation into 1L LB culture. The cells were induced at mid-log phase with the addition of 0.5mM IPTG (isopropyl-β-D-thiogalactopyranoside) and grown overnight at 16°C and harvested. Cells were lysed and centrifuged at 4000rpm for 30 minutes in buffer A (20mM Tris pH 8.0, 500mM NaCl, 5mM β-mercaptoethanol and 5% glycerol) sonicated 10 times on pulse for 20 s with 40 s rest between the pulses. The cell lysate was centrifuged at 16000 rpm for 1 hour at 4°C and the resulting supernatant was passed through the pre-equilibrated Ni-NTA resin column. The protein bound resins were washed by 100mL buffer B (20mM Tris pH 8.0, 500mM NaCl, 5mM β-mercaptoethanol, 5% glycerol and 50mM imidazole) and eluted with buffer C (20mM Tris pH 8.0, 500mM NaCl, 5mM β-mercaptoethanol, 5% glycerol and 300mM imidazole). The ULP1 protease was added to the elutant to remove the sumo-tag from DiCas7-11 while dialyzing at 4°C overnight. The digested protein solution was diluted 2-fold before being loaded onto a heparin column pre-equilibrated with buffer D (20mM Tris pH 8.0, 250mM NaCl, 5mM β-mercaptoethanol and 5% glycerol). The bound protein was eluted with a salt gradient. Pooled fractions were further purified on a gel filtration column in buffer E (20mM Tris pH 8.0, 500mM NaCl, 2mM DTT and 5% glycerol). The protein containing fractions were pooled, concentrated to 21mg/mL, aliquoted and stored at −80 °C for future use. The DiCas7-11 mutants were prepared by Q5 2X master mix mutagenesis kit (New England Biolabs) using primers listed in Supplementary Table 2 and purified similarly as the wild-type DiCas7-11.

### In vitro transcription and purification

For synthesis of 59nt pre-crRNA, DNA oligonucleotides appended with T7 promoter sequence were ordered from Eurofins (Supplementary Table 2). The complementary oligos, 50μM in concentration, were annealed at 95°C followed by gradual cooling to 25°C at 1°C per minute rate. Next, 2.5μL of annealing reaction was mixed with a transcription reaction master-mix (50 mM Tris pH 8.0, 10 mM DTT 20 mM MgCl_2_, 0.5 mM NTPs, and 48 μg/mL T7 RNA polymerase in a 50μL reaction. The in-vitro transcription reaction was kept overnight at 37°C and treated with 2U of Turbo DNase (Invitrogen) for 1 hour at 37°C. The final product was purified by Monarch RNA Cleanup kit (New England Biolabs), eluted in water, flash frozen using liquid nitrogen and stored at −80 °C.

### RNA target cleavage

The in-vitro target RNA cleavage assays were performed in a cleavage buffer containing 40 mM Tris pH 8.0, 70mM sodium chloride, 10 mM MgCl_2_. A binary complex was first prepared by incubating 400nM Cas7-11 with 500nM crRNA for 30 min at 37 °C. The resulting samples were then incubated with 500nM target RNA (5Cy3 labelled or non-labelled) for 1h at 37 °C. The reactions were stopped by using 2x formamide dye (95% formamide, 0.025% SDS, 0.025% xylene cyanol FF, 0.5 mM EDTA). The samples were heated at 95 °C for 5 min and separated by 8 M Urea, 12% polyacrylamide-gel electrophoresis (PAGE) gels in 1x Tris Borate EDTA (TBE) running buffer and were visualized by staining with SYBR Gold II (Invitrogen) stain or fluorescence imager.

### Pre-crRNA processing

500nM of DiCas7-11 was incubated with 500nM of pre-crRNA at 37°C for 30 min in a 15 μL reactions containing 1X processing buffer (40 mM Tris, pH 8.0, 70 mM NaCl). The reactions were stopped by using 2x formamide dye (95% formamide, 0.025% SDS, 0.025% xylene cyanol FF, 0.5 mM EDTA). The samples were heated at 95 °C for 5 min and separated by 8 M Urea, 12% polyacrylamide-gel electrophoresis (PAGE) gels in 1x TBE running buffer and were visualized by staining with SYBR Gold II (Invitrogen) stain or fluorescence imager.

### Sample preparation and data collection for cryo-EM studies

To reconstitute the ternary complex, 400μg wild-type DiCas7-11 protein was incubated with 1.5 molar excess of a pre-crRNA in the buffer (30mM Tris pH 8.0, 60mM NaCl) at 37°C for 45 minutes followed by separation on a Superdex 200 increase 10/300 gel filtration column in the buffer (30mM HEPES pH 7.5, 180mM NaCl, 10mM MgCl_2_ and 2mM TCEP). The peak fraction at 0.3mg/mL determined by UV 280nm absorbance was collected and further incubated with 2 molar excess target RNA at 37°C for 10 minutes before grid preparation. The Cas7-11-crRNA-target-RNA sample in 5μL volume was added onto glow-discharged 300 mesh Cu R1.2/1.3 holey carbon grids (Quantifoil) with extra layer of carbon (2 nm), blotted for 3s at 100% humidity using FEI Vitrobot Mark IV. After flash-freezing in liquid ethane, the grids were transferred to liquid nitrogen for storage until cryo-EM imaging.

The Cas7-11 ternary complex micrographs were collected using EPU software on the Krios G3i cryo TEM (ThermoFisher Scientific) equipped with Gatan Bioquantum K3 direct electron detector (Gatan) with 15 eV energy filter in a counted super-resolution mode. All 4177 images were collected at a dose rate of 60 e-/Å^2^ with 1e−/Å^2^ per frame in a pixel size of 0.825 Å/pixel. Motion correction was performed in bin 2 using MotionCorr 2 ^20^ in a wrapper provided in Relion 4.0 ^21^ and contrast transfer function (CTF) parameters were estimated with Gctf ^22^ implemented in cryo-SPARC ^23^. The stack was generated and imported to cryoSPARC for particle picking and 2D classification. The images with bad ice, astigmatism, drift, and poor sample quality were rejected resulting in 4168 images for further processing and particle picking, which resulted in a total of 1,301,452 particles. Several rounds of 2D classification led to 645,053 particles with good quality. RELION-4.0 was used to classify the particles, which led to further reduction of particles to 226,320 based on high-resolution features for reconstruction.

### Model building and refinement

The DiCas7-11 protein was built from an AlphaFold ^24^ predicted structure model using the program COOT ^25^. The final DiCas7-11-crRNA-target RNA complex was refined in PHENIX ^26^ to satisfactory stereochemistry and density correlation parameters (Supplementary Table 1).

## Data Availability

The atomic coordinates and associated density maps have deposited at Protein Data Bank with accession codes 8D1V & EMD-27138.

## Acknowledgments

This work was supported by NIH grant R01 GM101343 to H.L. The authors also acknowledge the use of instruments at the Biological Science Imaging Resource supported by Florida State University. The Titan was funded from NIH grant S10 RR025080. The BioQuantum/K3 was funded from NIH grant U24 GM116788. The Vitrobot Mk IV was funded from NIH grant S10 RR024564. The Solaris Plasma Cleaner was funded from NIH grant S10 RR024564. The DE-64 was funded from NIH grant U24 GM116788. The Laboratory for BioMolecular Structure (LBMS) is supported by the DOE Office of Biological and Environmental Research (KP160711).

## Author Contributions

H.G. A.D. and H.L. designed the experiments. H.G. purified the samples with the assistance of A.D. H.G. and J.R. prepared cryoEM grids and collected data. J.R. did cryoEM analysis with the assistance of H.G. H.G. J. R. and H.L analyzed data, wrote and edited manuscript and figures with the assistance of A.D.

## Supplementary Materials

**Supplementary Figure 1.**
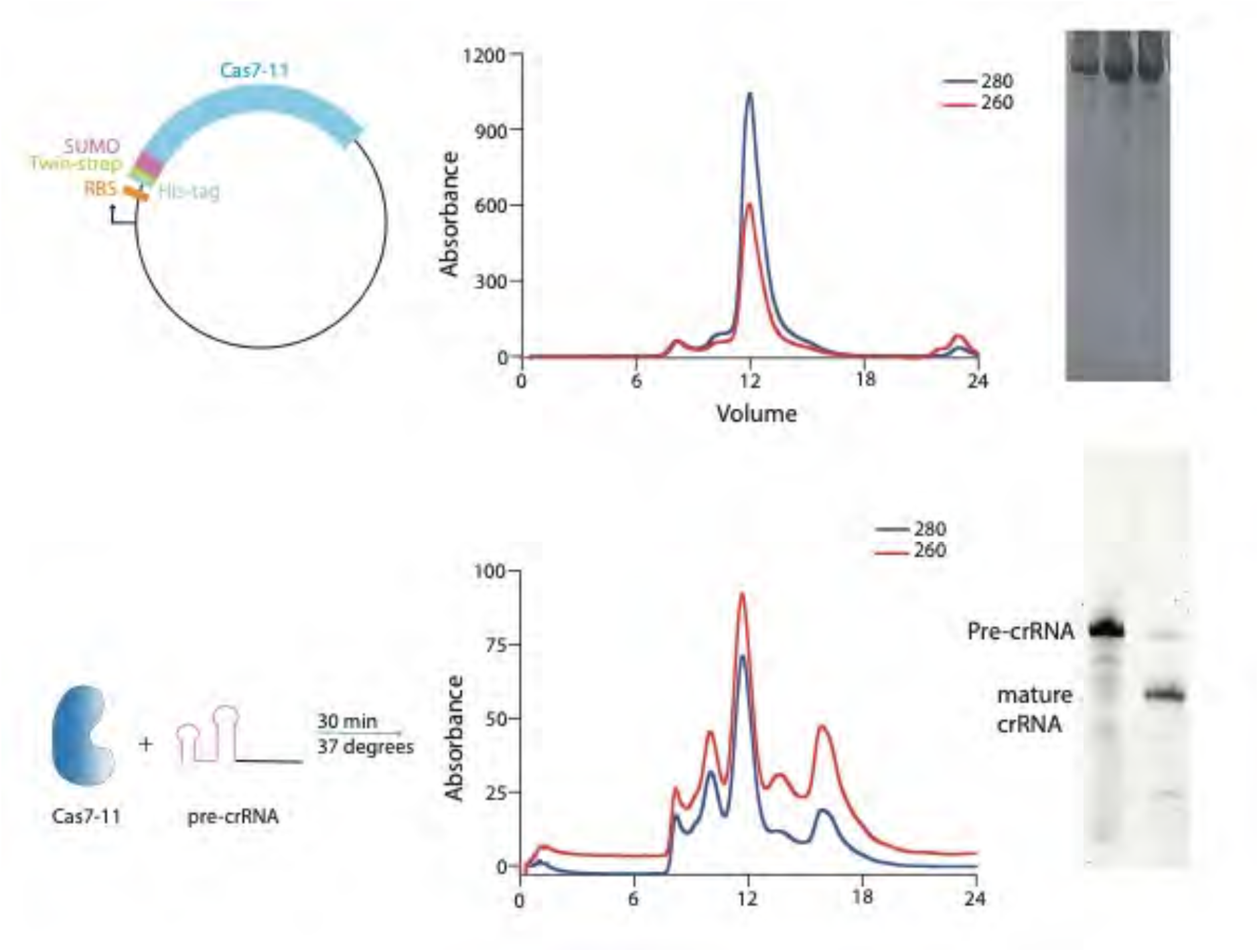
Top, schematic of the vector used for over-expression of apo DiCas7-11, the elution profile of DiCas7-11 on size exclusion chromatography, and gel analysis of the purified protein. Bottom, schematic of DiCas7-11 and pre-crRNA incubation for binary complex formation, elution profile of DiCas7-11-crRNA binary complex, and gel analysis of the crRNA in the peak fraction.

**Supplementary Figure 2.**
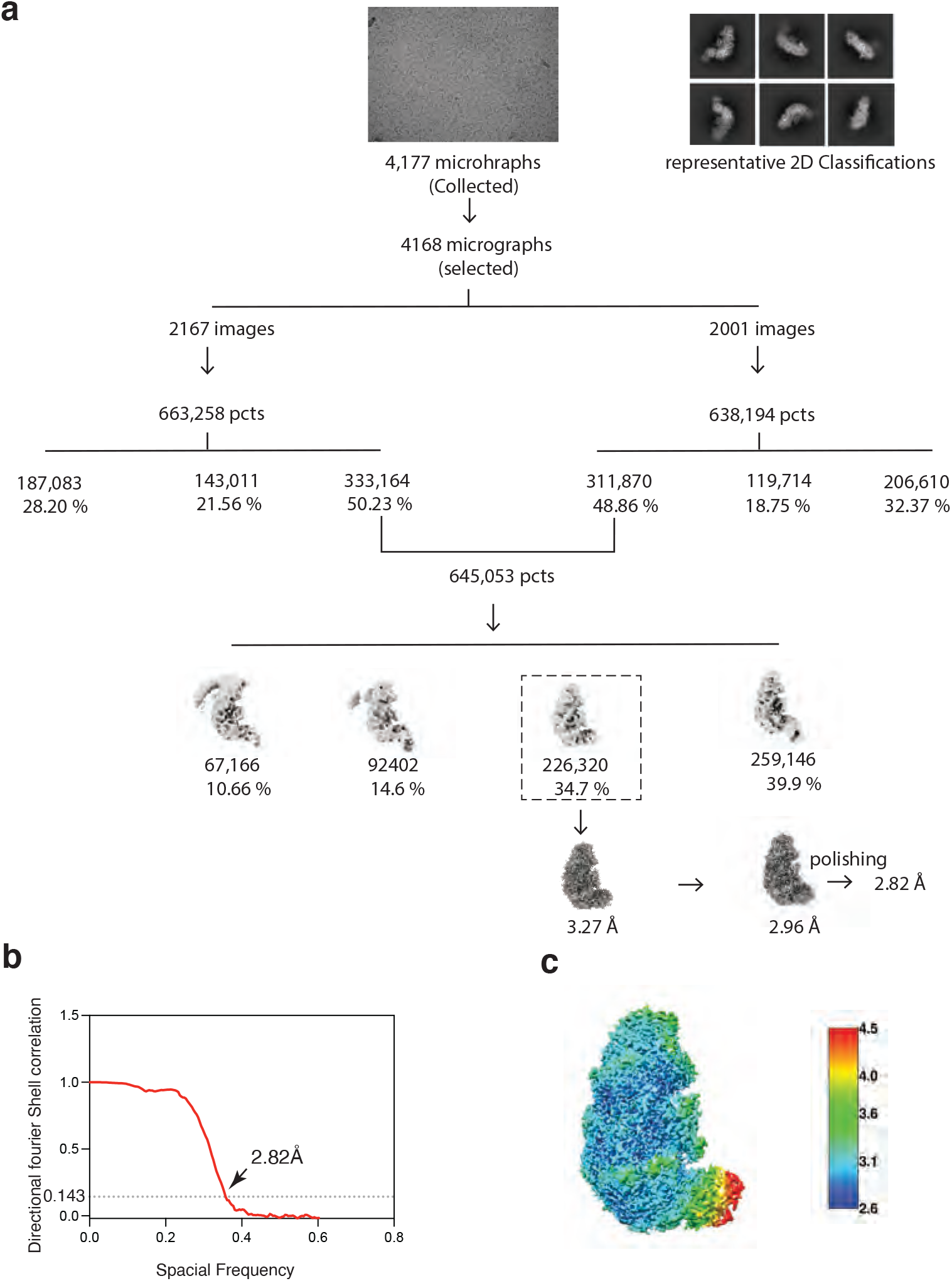
(a) Work-flow of the cryo-EM image processing and 3D reconstruction of the DiCas7-11 ternary complex. The numbers indicate the number of particles while the percent indicates the percent to the total particles for each class. (b) Fourier Shell Correlations (FSC) of DiCas7-11-crRNA-ternary complex reconstruction, with the FSC cutoff 0.143, marked with a gray line and final resolution. (c) The local resolution of the final density map obtained from ResMap is colored as the color scale bar.

**Supplementary Figure 3.**
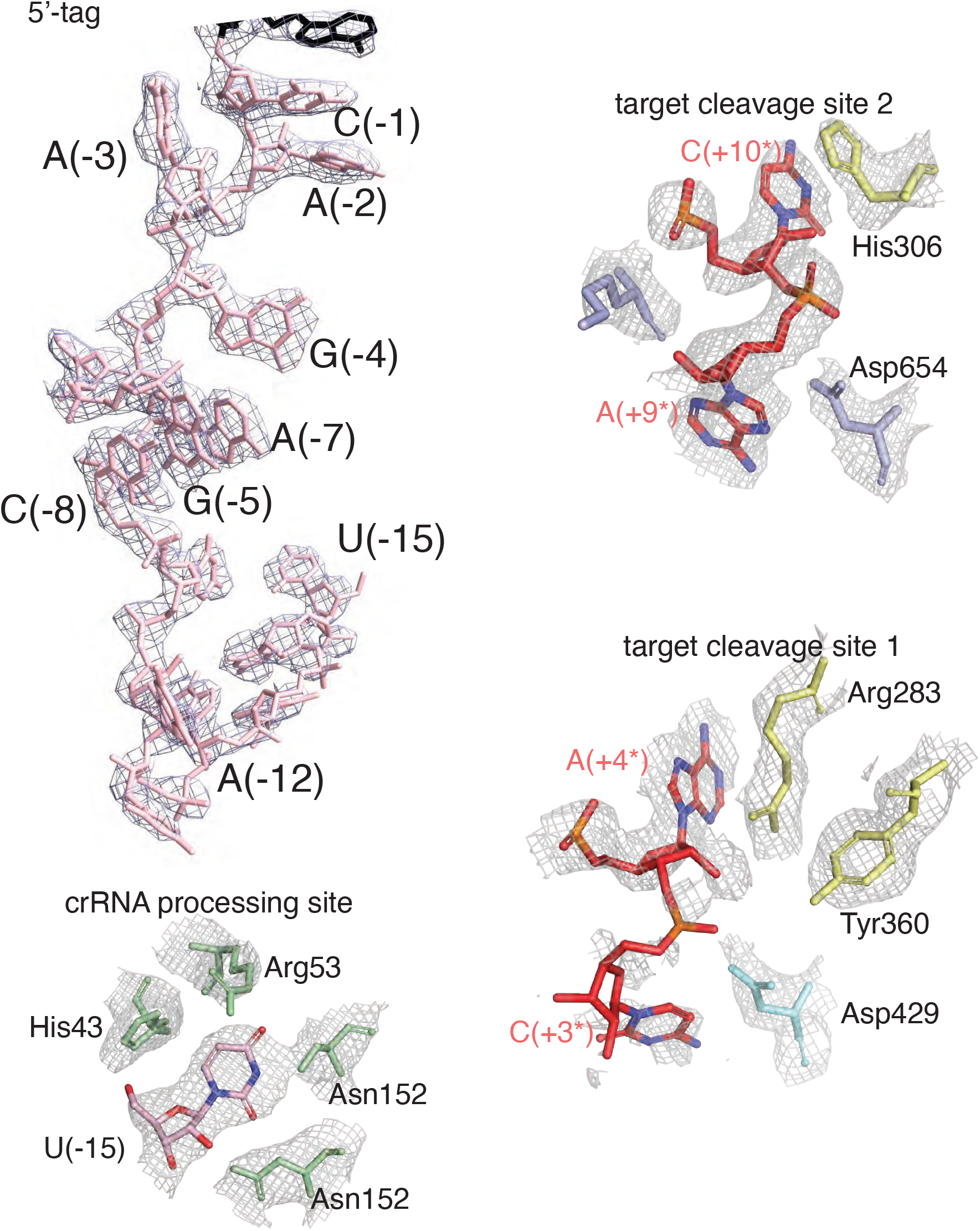
Electron potential density maps for selected regions where nucleotide positions are labeled: around the 5’ tag region, the crRNA processing site, and the two target cleavage sites.

**Supplementary Figure 4.**
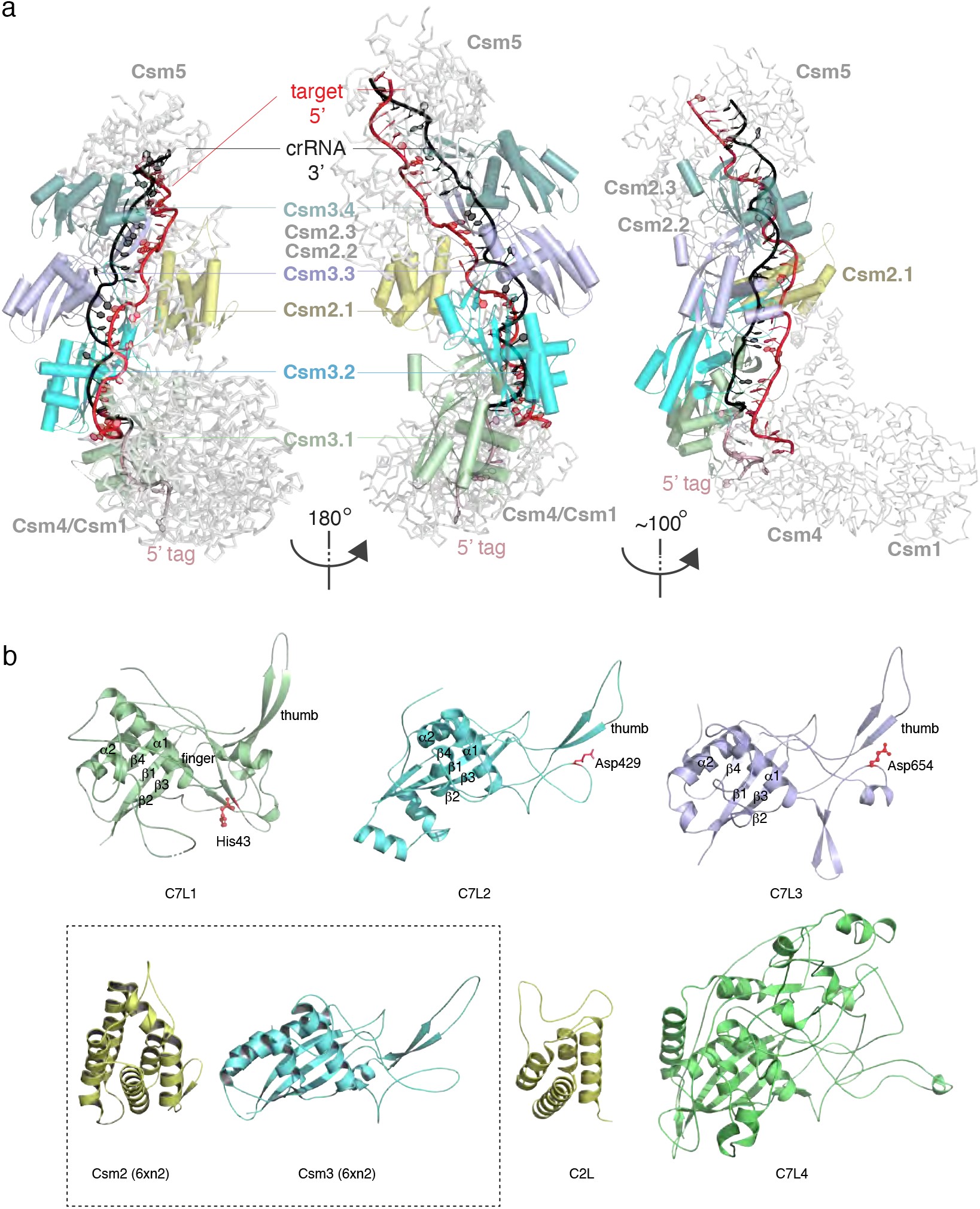
Comparison of DiCas7-11 to the homologous multi-subunit Csm complex from *Lactococcus lactis* (LlCsm) (a) and its individual protein domains (b). (a). LlCsm overview in three orientations in carton representations. The two left views are oriented exactly the same as those of DiCas7-11 shown in Figure 1. Subunits that superimposed with DiCas7-11 domains and RNA are labeled and colored identically. (b). The four Cas7-like (C7L) domains are superimposed and shown separately in cartoon representations and colored as those in Figure 1. The Cas11-like (C11L) is shown in cartoon representations. Key elements discussed in the text are labels. Boxed structures are the corresponding subunits for C11L and C7L, respectively, from LlCsm (PDB id: 6xn2).

**Supplementary Figure 5.**
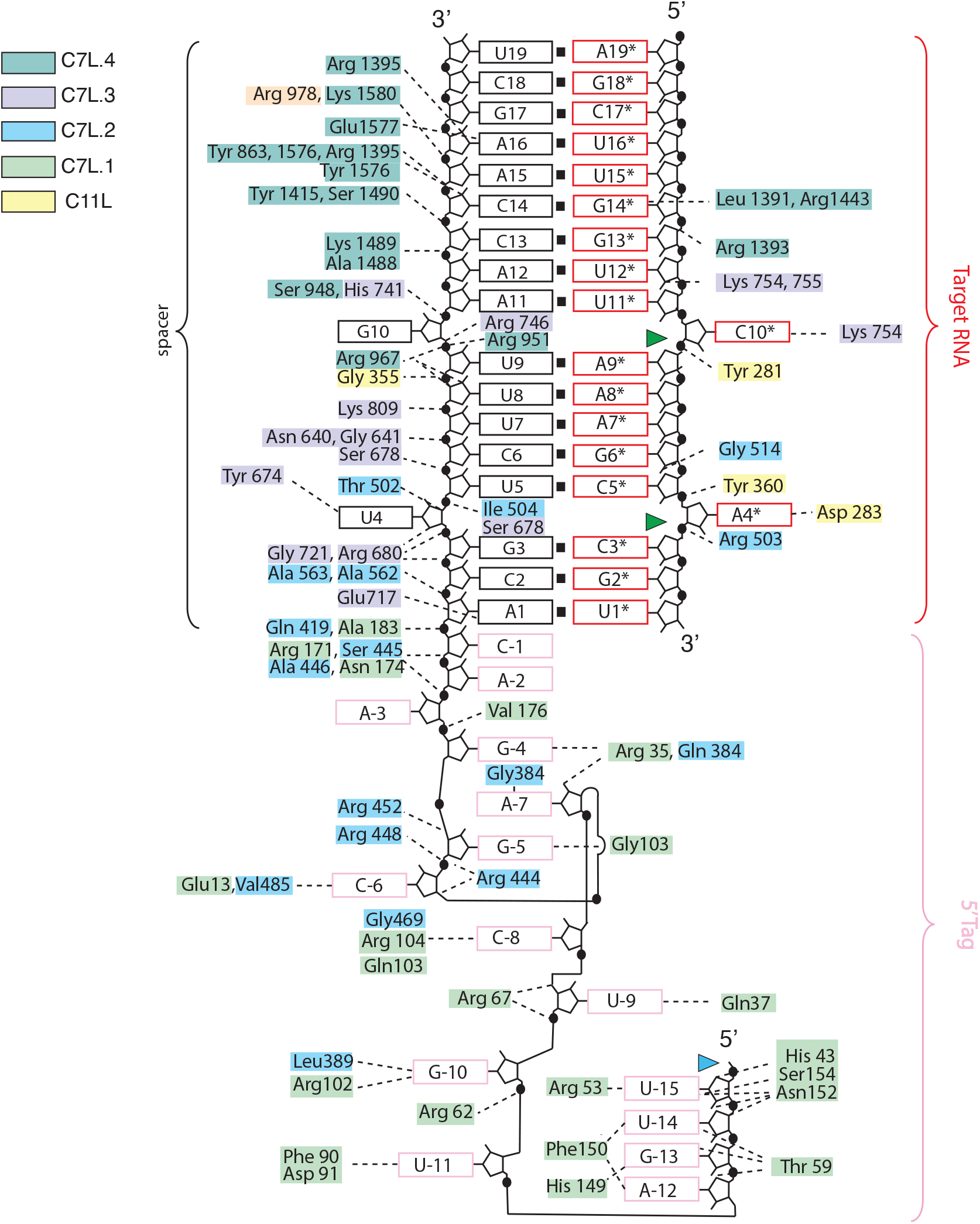
Schematic RNA-protein interactions observed in the DiCas7-11-crRNA-target RNA ternary complex structure. Protein residues are colored according to the scheme used in Figure 1 that corresponding to respective domains. The black, red and light pink color boxes denote spacer, repeat and target RNA. The green and blue colored triangles indicate target RNA and pre-crRNA processing sites, respectively.

**Supplementary Figure 6.**
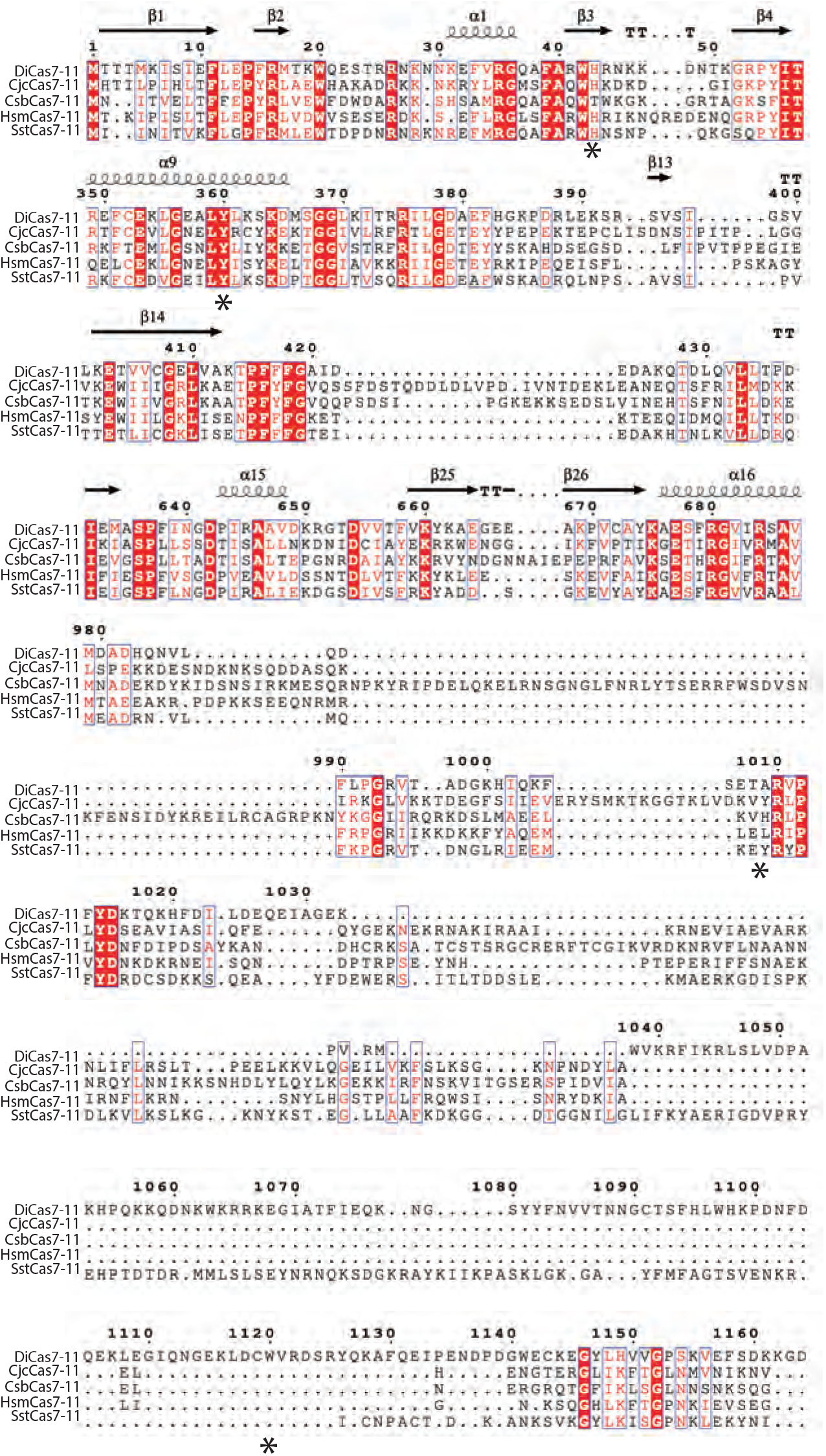
Sequence comparison among Cas7-11 proteins overlays with the secondary structure structures of DiCas7. Regions of interests and described in the main text are marked by asterisks. DiCas7-11 is from *Desulfonema ishimotonii*; CjcCas7-11 is from *Candidatus Jettenia caeni;* CsbCas7-11 is from *Candidatus Scalindua broadae;* HsmCas7-11 and SstCas7-11 source organism is unknown.

**Supplementary Table 1.**
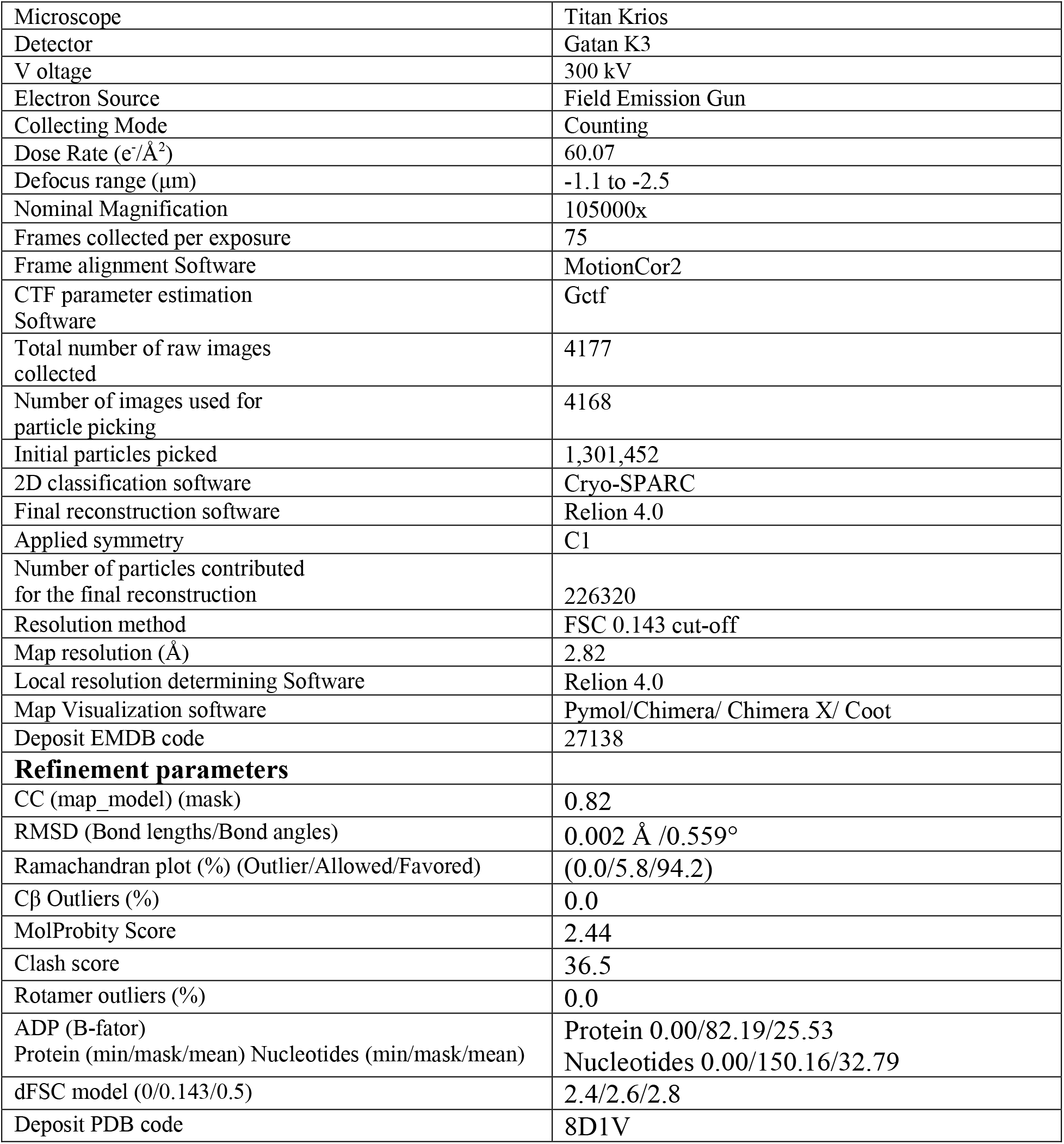
Data acquisition and processing parameters. Related to Figures 2-4

**Supplementary Table 2:**
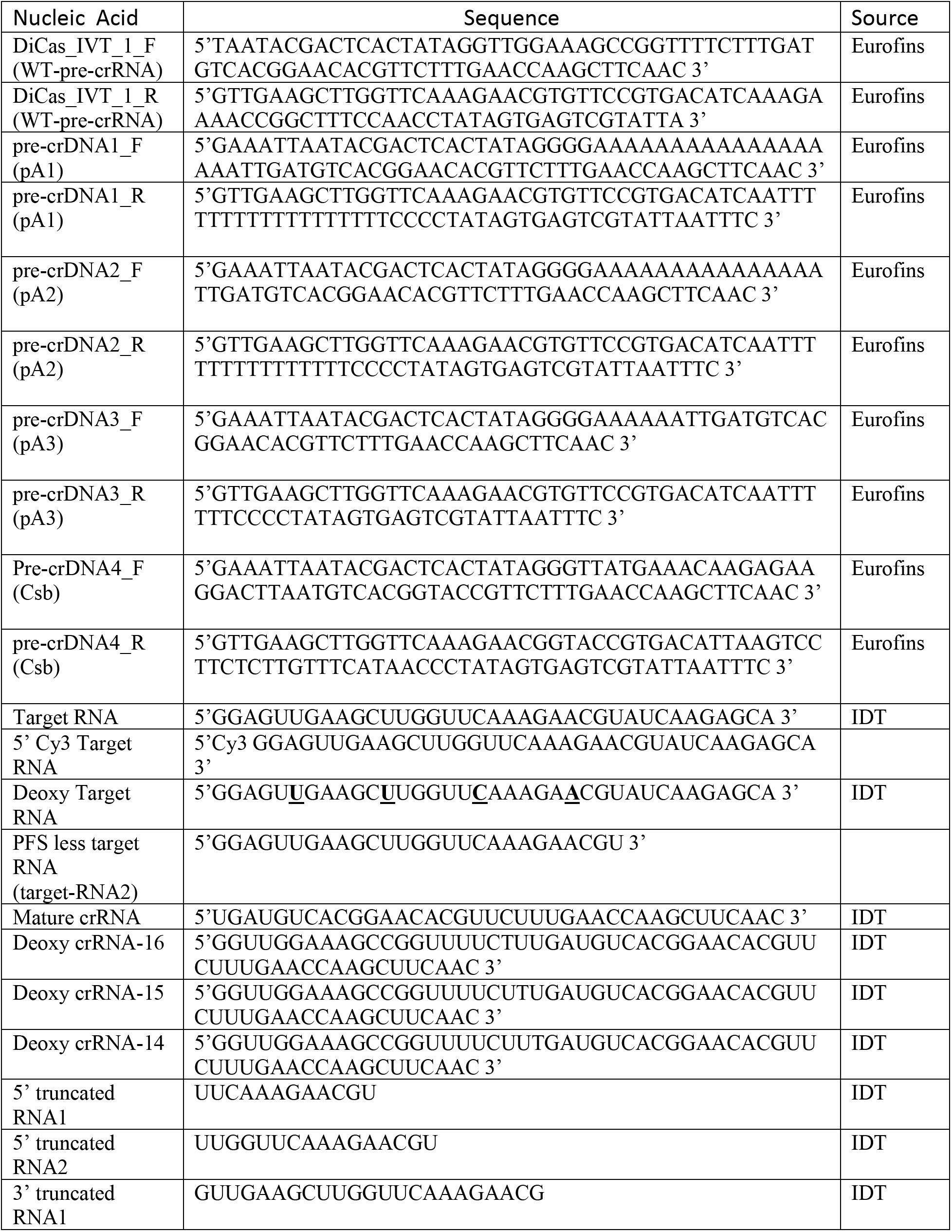
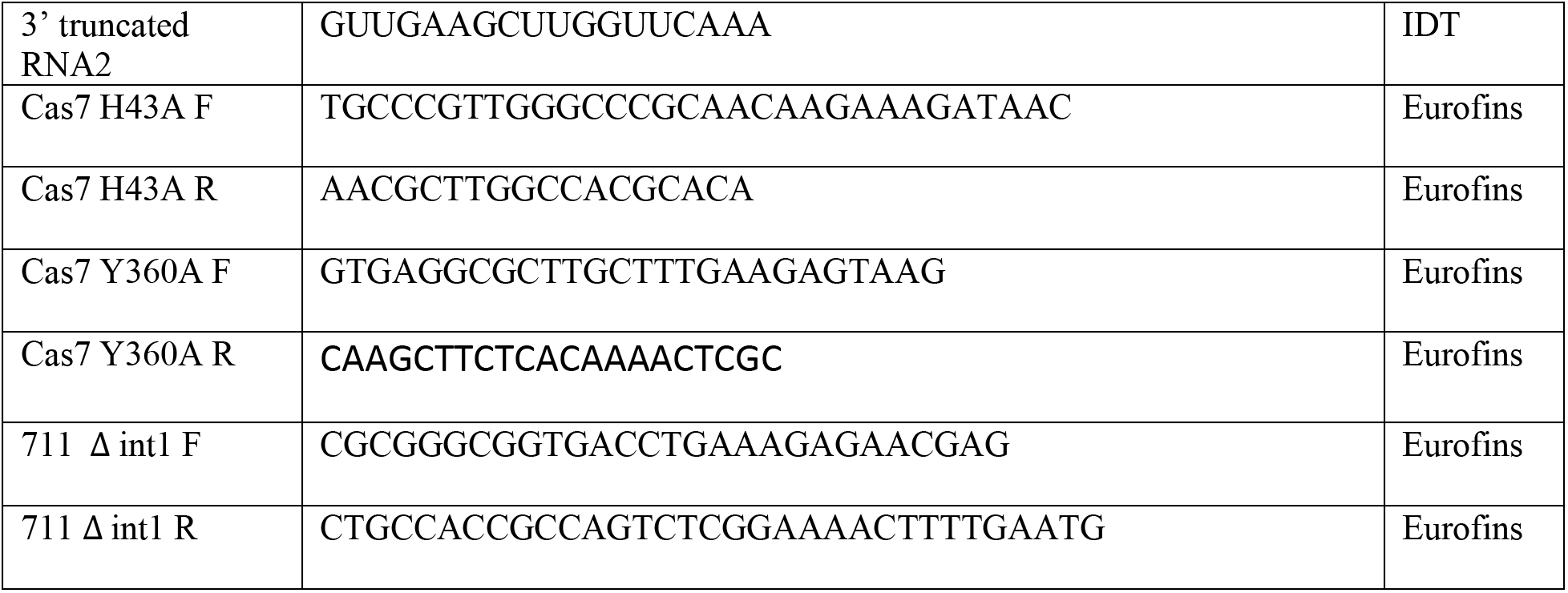
Ribonucleic acid sequences used in this study. Related to Figures 1-4

## Full wwPDB EM Validation Report

### 1 Overall quality at a glance

The following experimental techniques were used to determine the structure: *ELECTRON MICROSCOPY*

The reported resolution of this entry is 2.82 Å.

Percentile scores (ranging between 0-100) for global validation metrics of the entry are shown in the following graphic. The table shows the number of entries on which the scores are based.

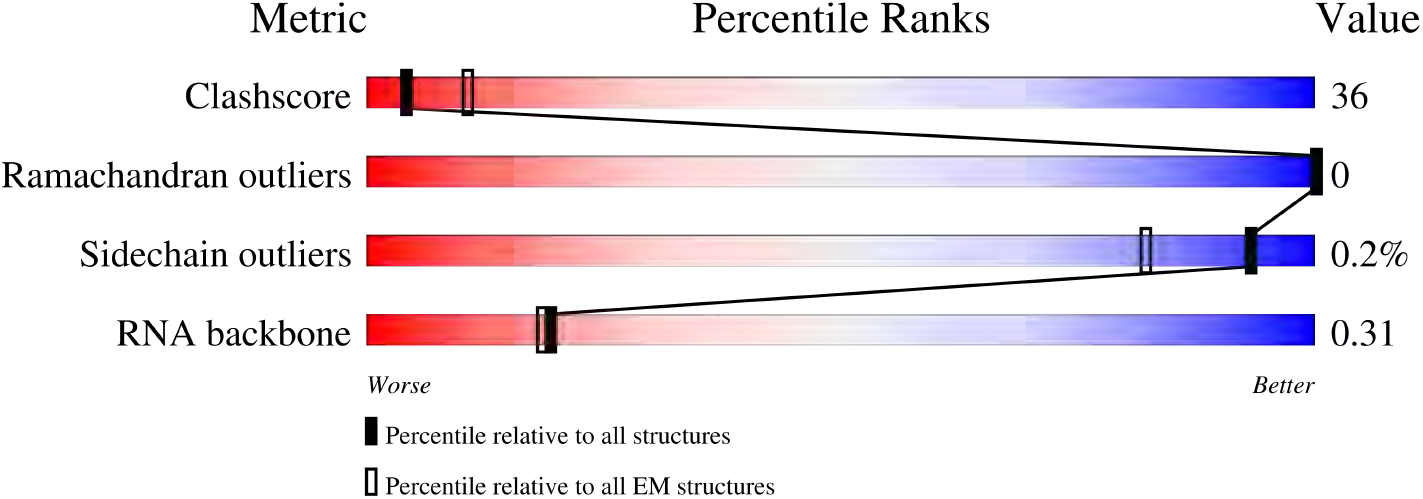

| Metric | Whole archive<br>(#Entries) | EM structures<br>(#Entries) |
| --- | --- | --- |
| Clashscore | 158937 | 4297 |
| Ramachandran outliers | 154571 | 4023 |
| Sidechain outliers | 154315 | 3826 |
| RNA backbone | 4643 | 859 |

The table below summarises the geometric issues observed across the polymeric chains and their fit to the map. The red, orange, yellow and green segments of the bar indicate the fraction of residues that contain outliers for *>*=3, 2, 1 and 0 types of geometric quality criteria respectively. A grey segment represents the fraction of residues that are not modelled. The numeric value for each fraction is indicated below the corresponding segment, with a dot representing fractions *<*=5% The upper red bar (where present) indicates the fraction of residues that have poor fit to the EM map (all-atom inclusion *<* 40%). The numeric value is given above the bar.

| Mol | Chain | Length | Quality of chain |
| --- | --- | --- | --- |
| 1 | A | 1253 | 20%<br>47%<br>53% |
| 2 | N | 34 | 9%<br>9%<br>59%<br>32% |
| 3 | J | 18 | 33%<br>11%<br>28%<br>61% |

### 2 Entry composition

There are 3 unique types of molecules in this entry. The entry contains 11148 atoms, of which 9 are hydrogens and 0 are deuteriums.

In the tables below, the AltConf column contains the number of residues with at least one atom in alternate conformation and the Trace column contains the number of residues modelled with at most 2 atoms.

- Molecule 1 is a protein called Di Cas7-11.

- Molecule 2 is a RNA chain called CRISPR RNA (34-MER).

| Mol | Chain | Residues | Atoms |  |  |  |  |  | AltConf | Trace |
| --- | --- | --- | --- | --- | --- | --- | --- | --- | --- | --- |
|  |  |  | Total | C | H | N | O | S |  |  |
| 1 | A | 1251 | 10045 | 6340 | 9 | 1797 | 1852 | 47 | 0 | 0 |

- Molecule 3 is a RNA chain called SS target RNA (5’-R(P*AP*GP*CP*UP*UP*GP*GP*U P*UP*CP*AP*AP*AP*GP*AP*AP*CP*G)-3’).

- Molecule 2 is a RNA chain called CRISPR RNA (34-MER).
| Mol | Chain | Residues | Atoms |  |  |  |  | AltConf | Trace |
| --- | --- | --- | --- | --- | --- | --- | --- | --- | --- |
|  |  |  | Total | C | N | O | P |  |  |
| 2 | N | 34 | 716 | 322 | 124 | 237 | 33 | 0 | 0 |

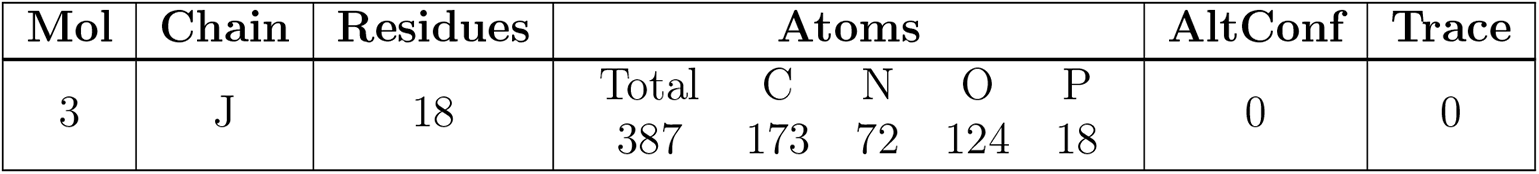

### 3 Residue-property plots

These plots are drawn for all protein, RNA, DNA and oligosaccharide chains in the entry. The first graphic for a chain summarises the proportions of the various outlier classes displayed in the second graphic. The second graphic shows the sequence view annotated by issues in geometry and atom inclusion in map density. Residues are color-coded according to the number of geometric quality criteria for which they contain at least one outlier: green = 0, yellow = 1, orange = 2 and red = 3 or more. A red diamond above a residue indicates a poor fit to the EM map for this residue (all-atom inclusion *<* 40%). Stretches of 2 or more consecutive residues without any outlier are shown as a green connector. Residues present in the sample, but not in the model, are shown in grey.

- Molecule 1: Di Cas7-11

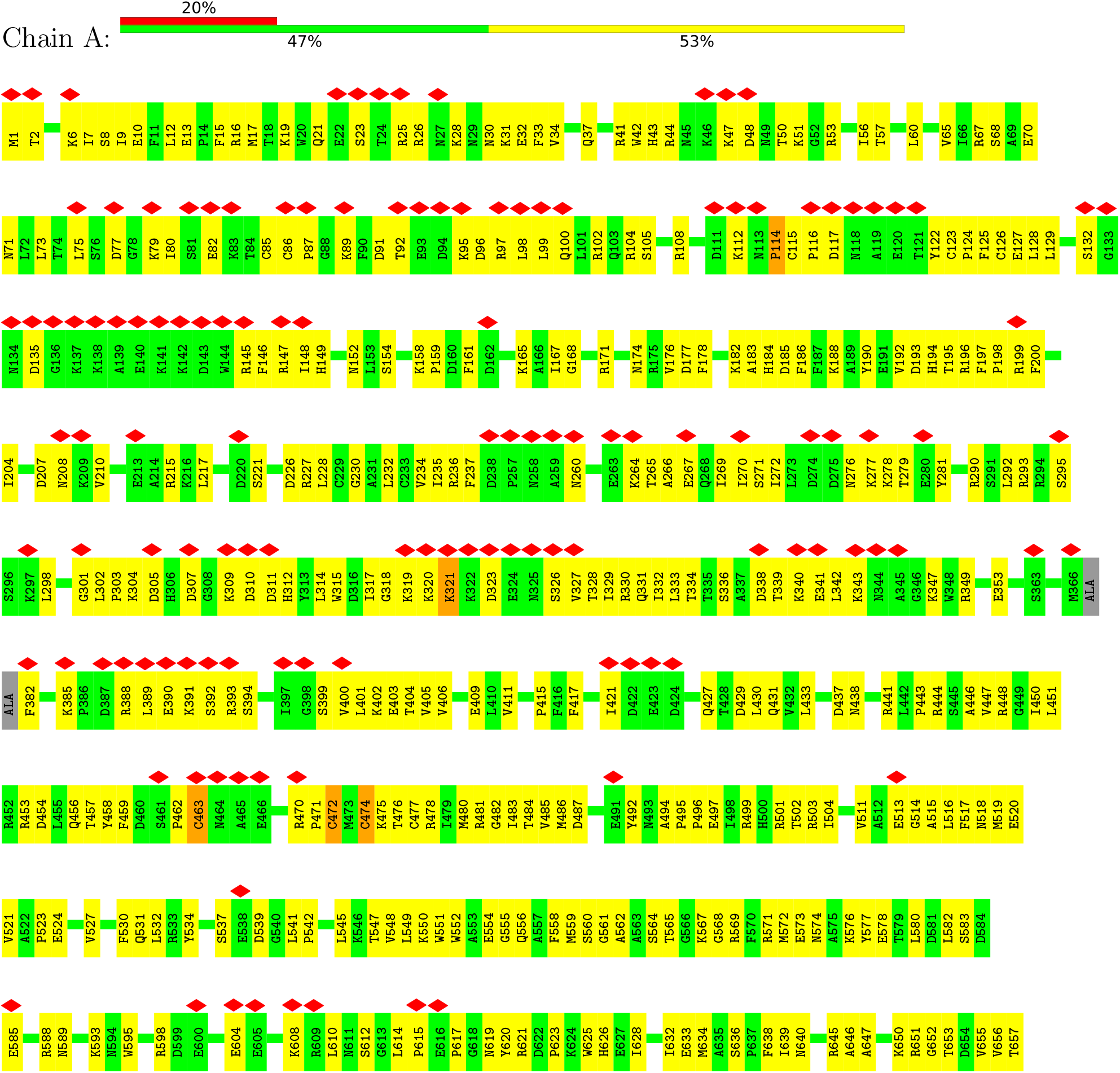
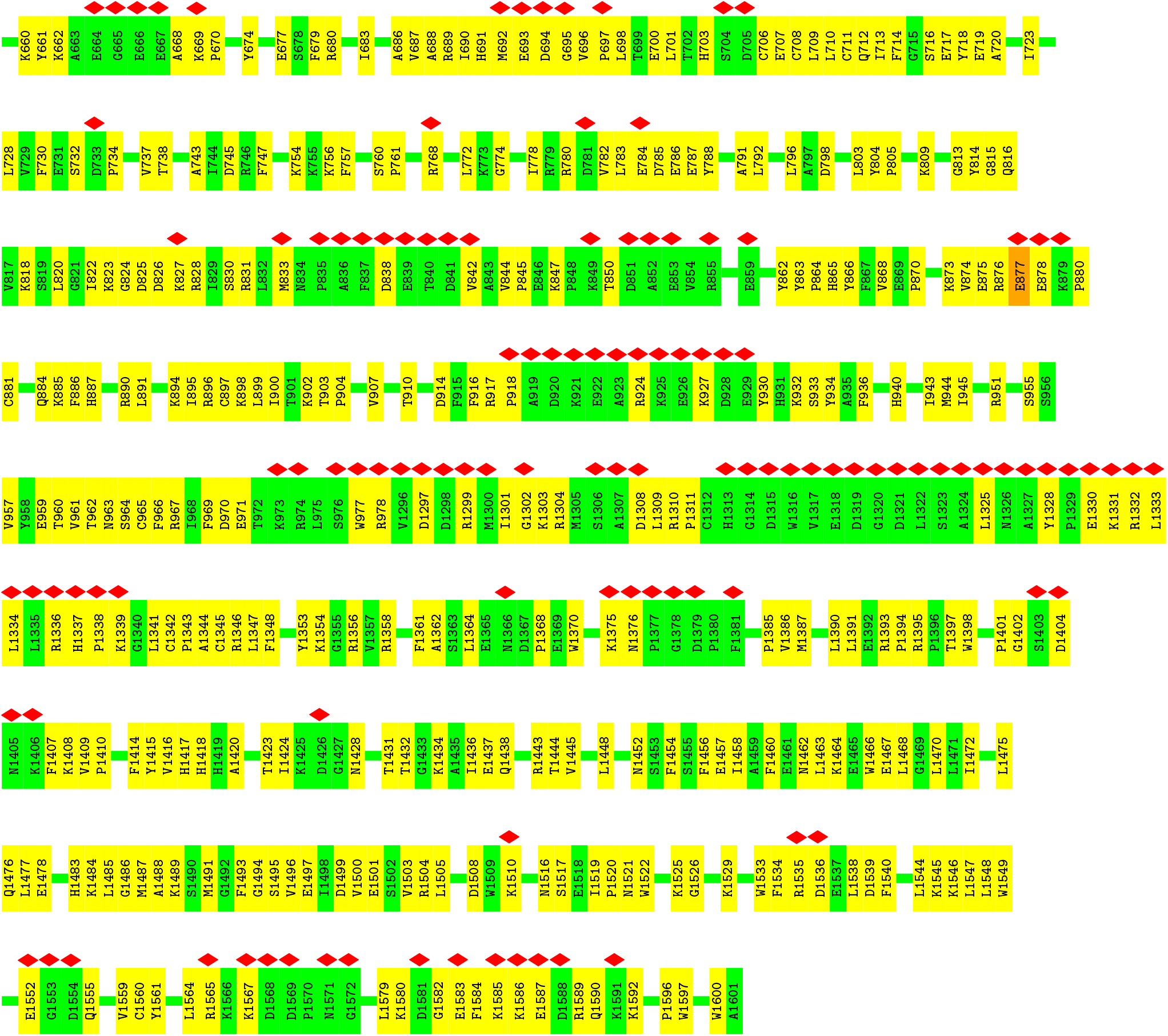
- Molecule 2: CRISPR RNA (34-MER)

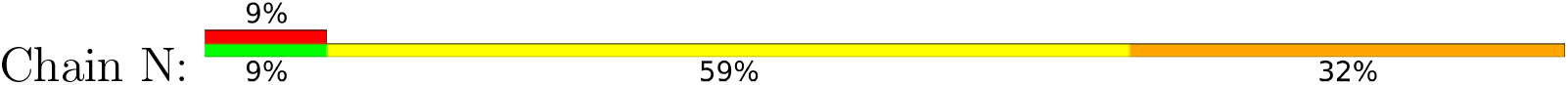
- Molecule 3: SS target RNA (5’-R(P*AP*GP*CP*UP*UP*GP*GP*UP*UP*CP*AP*AP*AP* GP*AP*AP*CP*G)-3’)

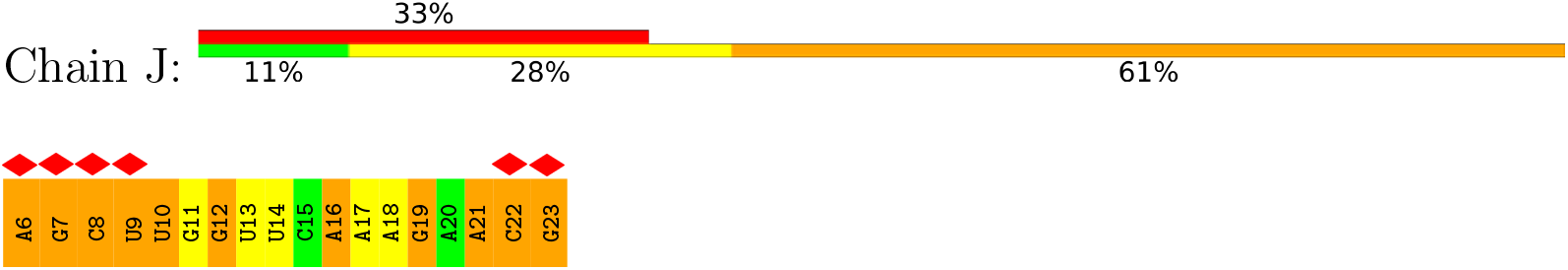

### 4 Experimental information

| Property | Value | Source |
| --- | --- | --- |
| EM reconstruction method | SINGLE PARTICLE | Depositor |
| Imposed symmetry | POINT, C1 | Depositor |
| Number of particles used | 226320 | Depositor |
| Resolution determination method | FSC 0.143 CUT-OFF | Depositor |
| CTF correction method | PHASE FLIPPING AND AMPLITUDE CORRECTION | Depositor |
| Microscope | FEI TITAN KRIOS | Depositor |
| Voltage (kV) | 300 | Depositor |
| Electron dose ($e^-/\text{\AA}^2$) | 60 | Depositor |
| Minimum defocus (nm) | 1000 | Depositor |
| Maximum defocus (nm) | 2200 | Depositor |
| Magnification | Not provided |  |
| Image detector | GATAN K3 BIOQUANTUM (6k x 4k) | Depositor |
| Maximum map value | 0.098 | Depositor |
| Minimum map value | -0.053 | Depositor |
| Average map value | 0.000 | Depositor |
| Map value standard deviation | 0.003 | Depositor |
| Recommended contour level | 0.0146 | Depositor |
| Map size (Å) | 249.0, 249.0, 249.0 | wwPDB |
| Map dimensions | 300, 300, 300 | wwPDB |
| Map angles (°) | 90.0, 90.0, 90.0 | wwPDB |
| Pixel spacing (Å) | 0.83, 0.83, 0.83 | Depositor |

### 5 Model quality

#### 5.1 Standard geometry

The Z score for a bond length (or angle) is the number of standard deviations the observed value is removed from the expected value. A bond length (or angle) with *|Z| >* 5 is considered an outlier worth inspection. RMSZ is the root-mean-square of all Z scores of the bond lengths (or angles).

| Mol | Chain | Bond lengths |  | Bond angles |  |
| --- | --- | --- | --- | --- | --- |
|  |  | RMSZ | #\|Z\| >5 | RMSZ | #\|Z\| >5 |
| 1 | A | 0.25 | 0/10270 | 0.51 | 6/13829 (0.0%) |
| 2 | N | 0.18 | 0/799 | 0.73 | 0/1242 |
| 3 | J | 0.17 | 0/433 | 0.71 | 0/673 |
| All | All | 0.25 | 0/11502 | 0.54 | 6/15744 (0.0%) |

There are no bond length outliers.

All (6) bond angle outliers are listed below:

There are no bond length outliers. All (6) bond angle outliers are listed below:
| Mol | Chain | Res | Type | Atoms | Z | Observed(°) | Ideal(°) |
| --- | --- | --- | --- | --- | --- | --- | --- |
| 1 | A | 472 | CYS | CA-CB-SG | 8.46 | 129.23 | 114.00 |
| 1 | A | 463 | CYS | CA-CB-SG | 7.29 | 127.13 | 114.00 |
| 1 | A | 474 | CYS | CA-CB-SG | 7.25 | 127.04 | 114.00 |
| 1 | A | 114 | PRO | CA-N-CD | -6.73 | 102.08 | 111.50 |
| 1 | A | 114 | PRO | N-CD-CG | -5.86 | 94.41 | 103.20 |
| 1 | A | 877 | GLU | C-N-CA | 5.36 | 135.10 | 121.70 |

There are no chirality outliers.

There are no planarity outliers.

#### 5.2 Too-close contacts

In the following table, the Non-H and H(model) columns list the number of non-hydrogen atoms and hydrogen atoms in the chain respectively. The H(added) column lists the number of hydrogen atoms added and optimized by MolProbity. The Clashes column lists the number of clashes within the asymmetric unit, whereas Symm-Clashes lists symmetry-related clashes.

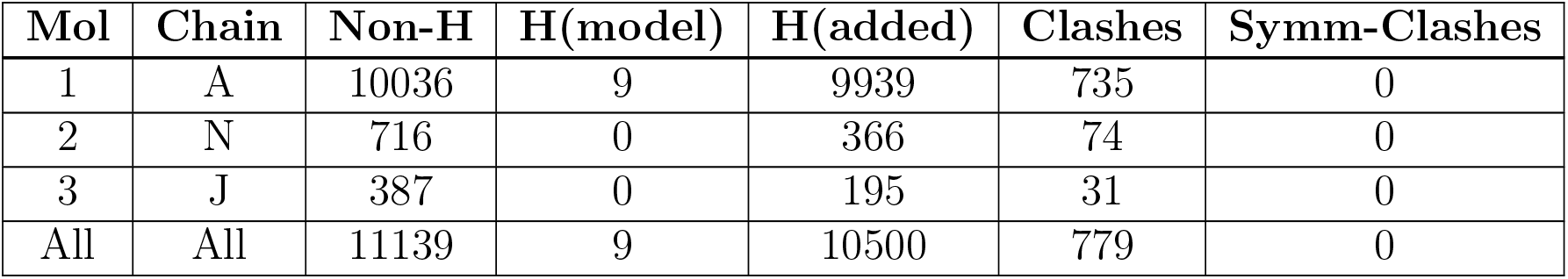

The all-atom clashscore is defined as the number of clashes found per 1000 atoms (including hydrogen atoms). The all-atom clashscore for this structure is 36.

All (779) close contacts within the same asymmetric unit are listed below, sorted by their clash magnitude.

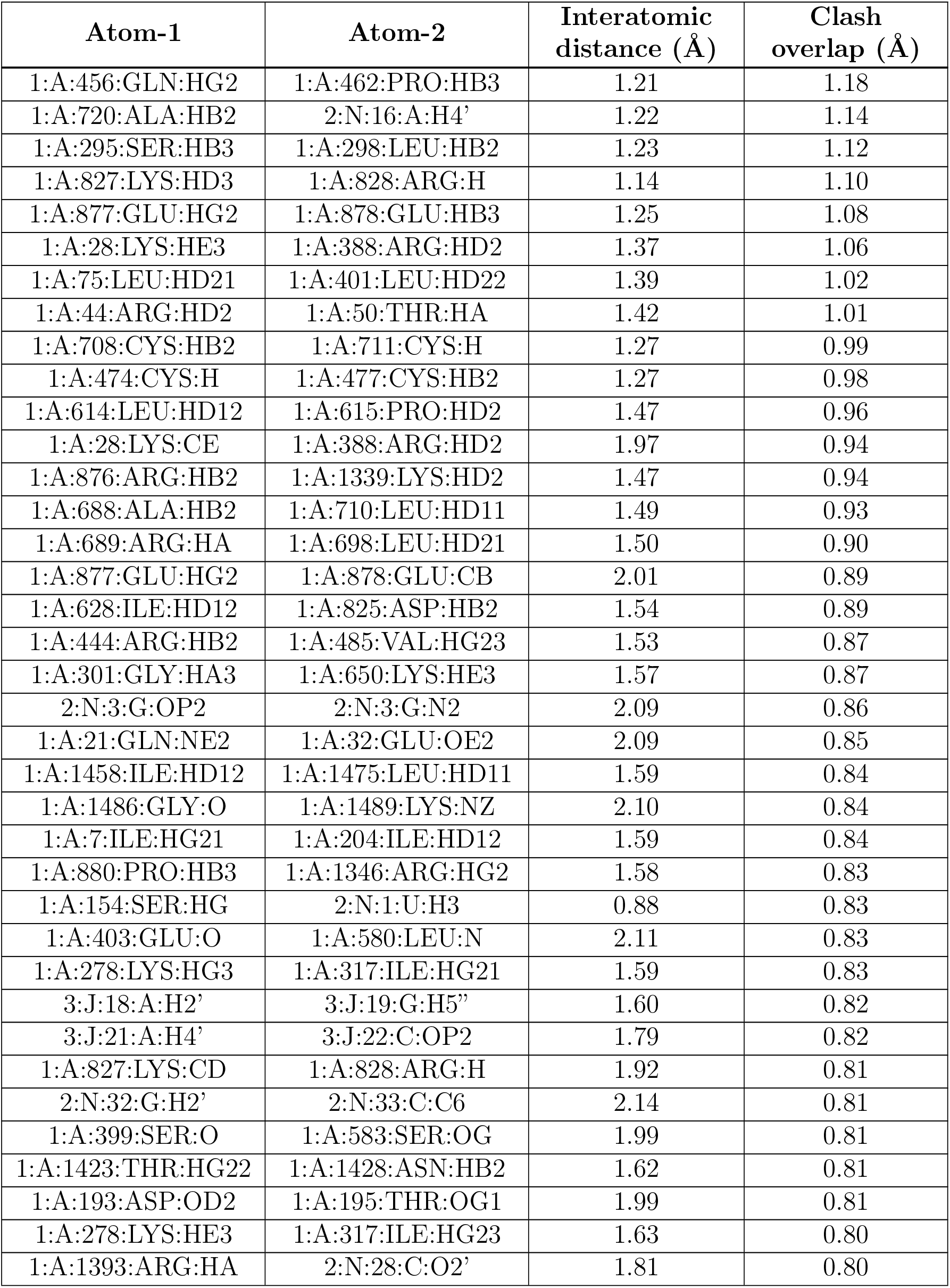
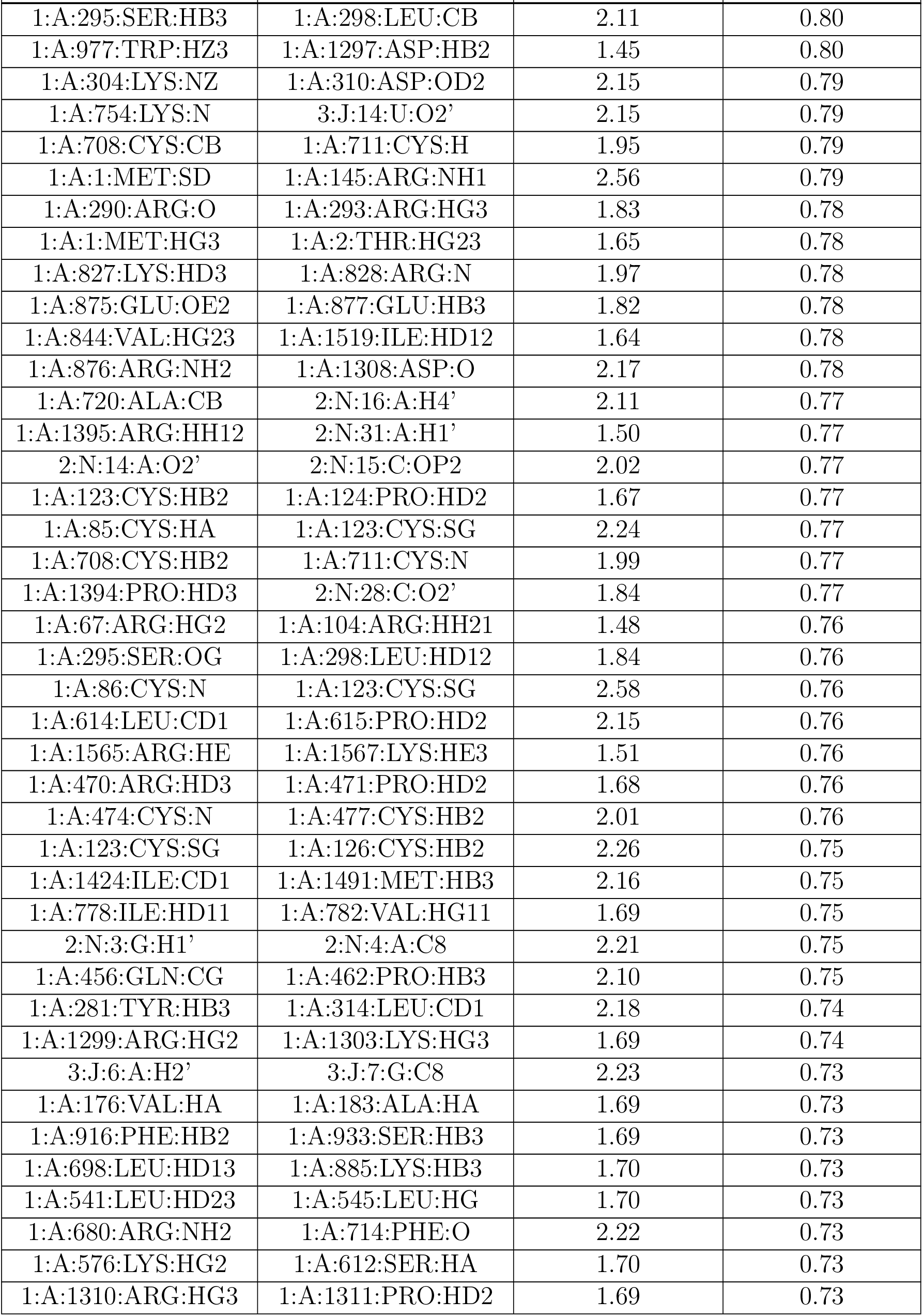
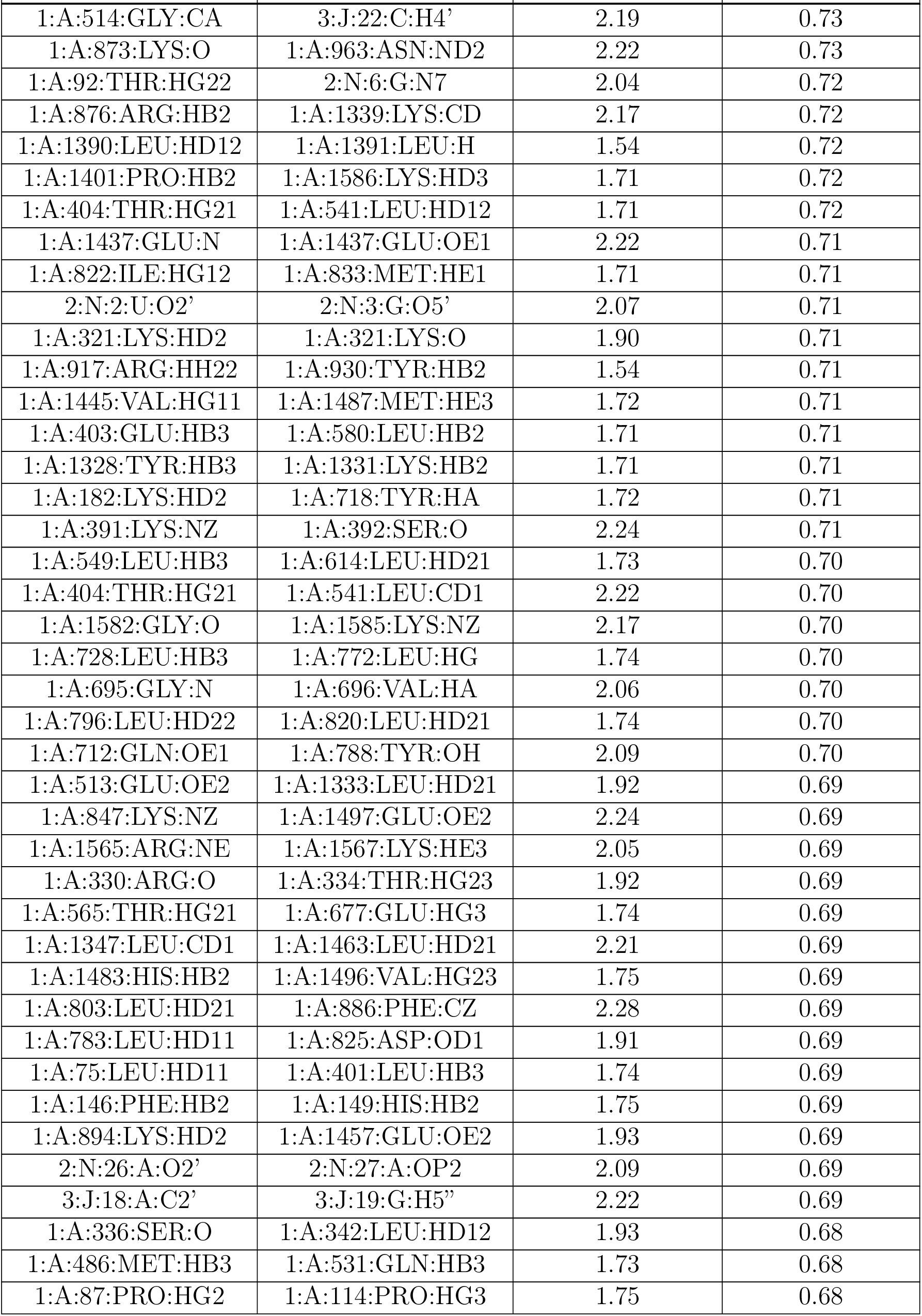
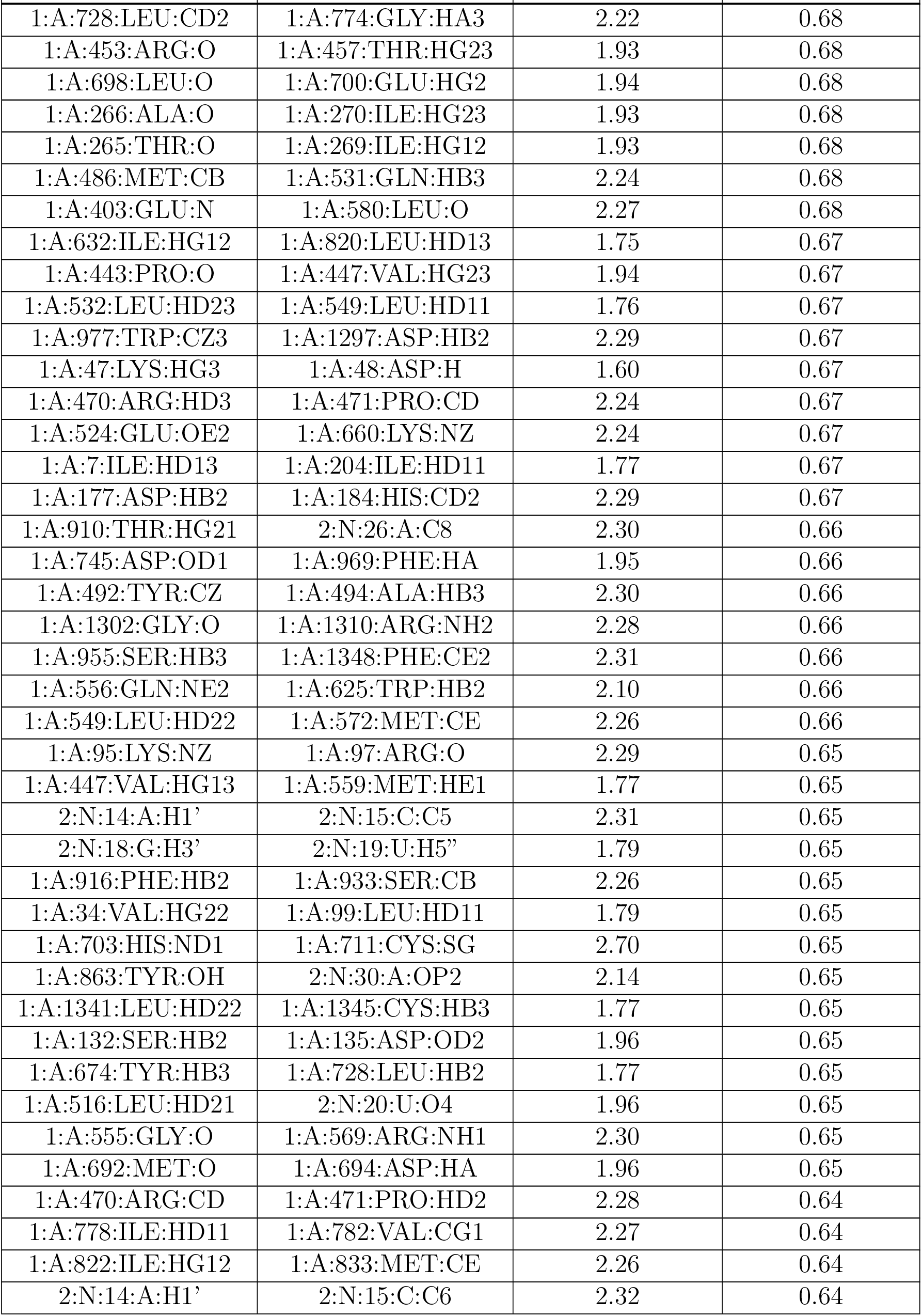
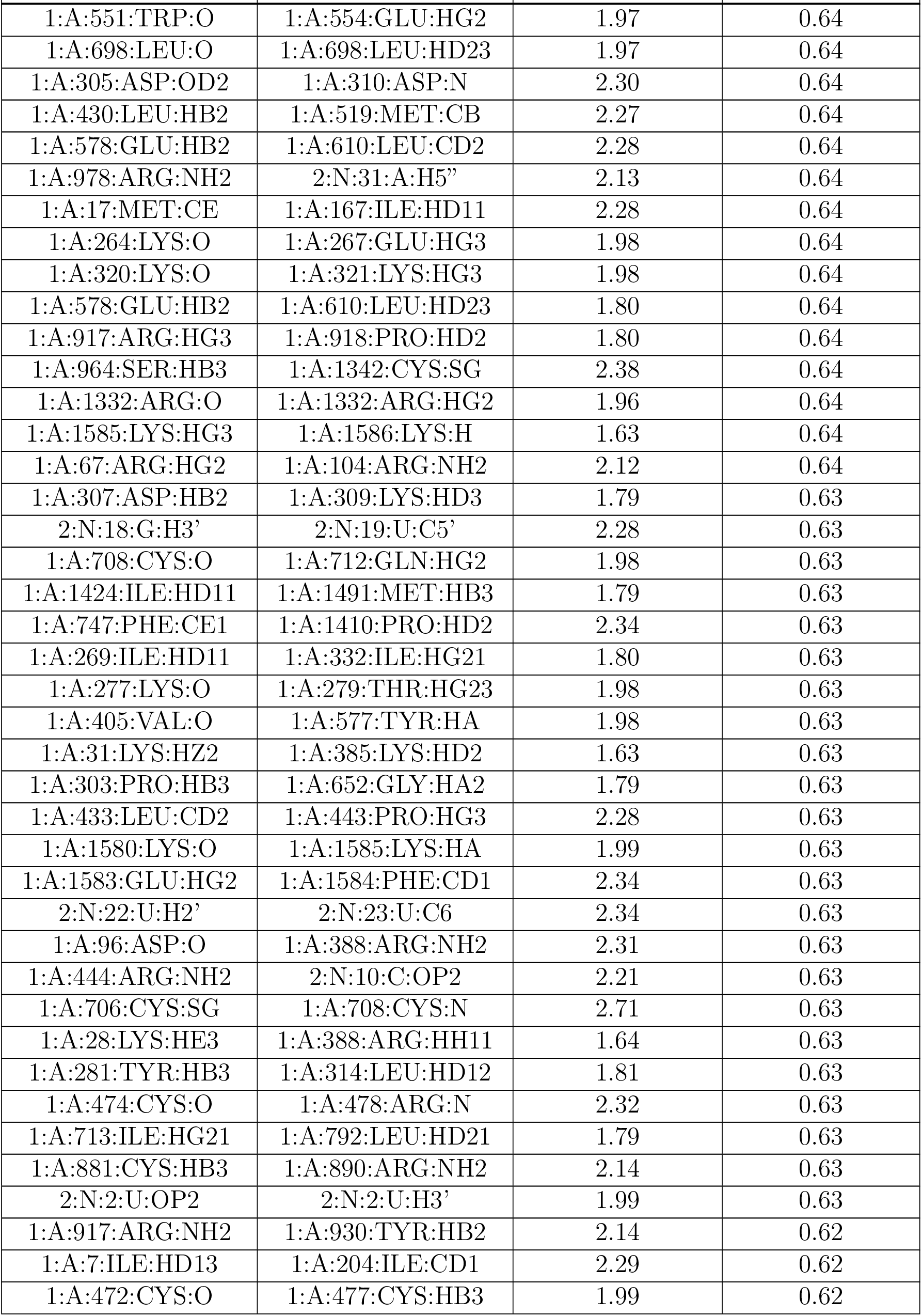
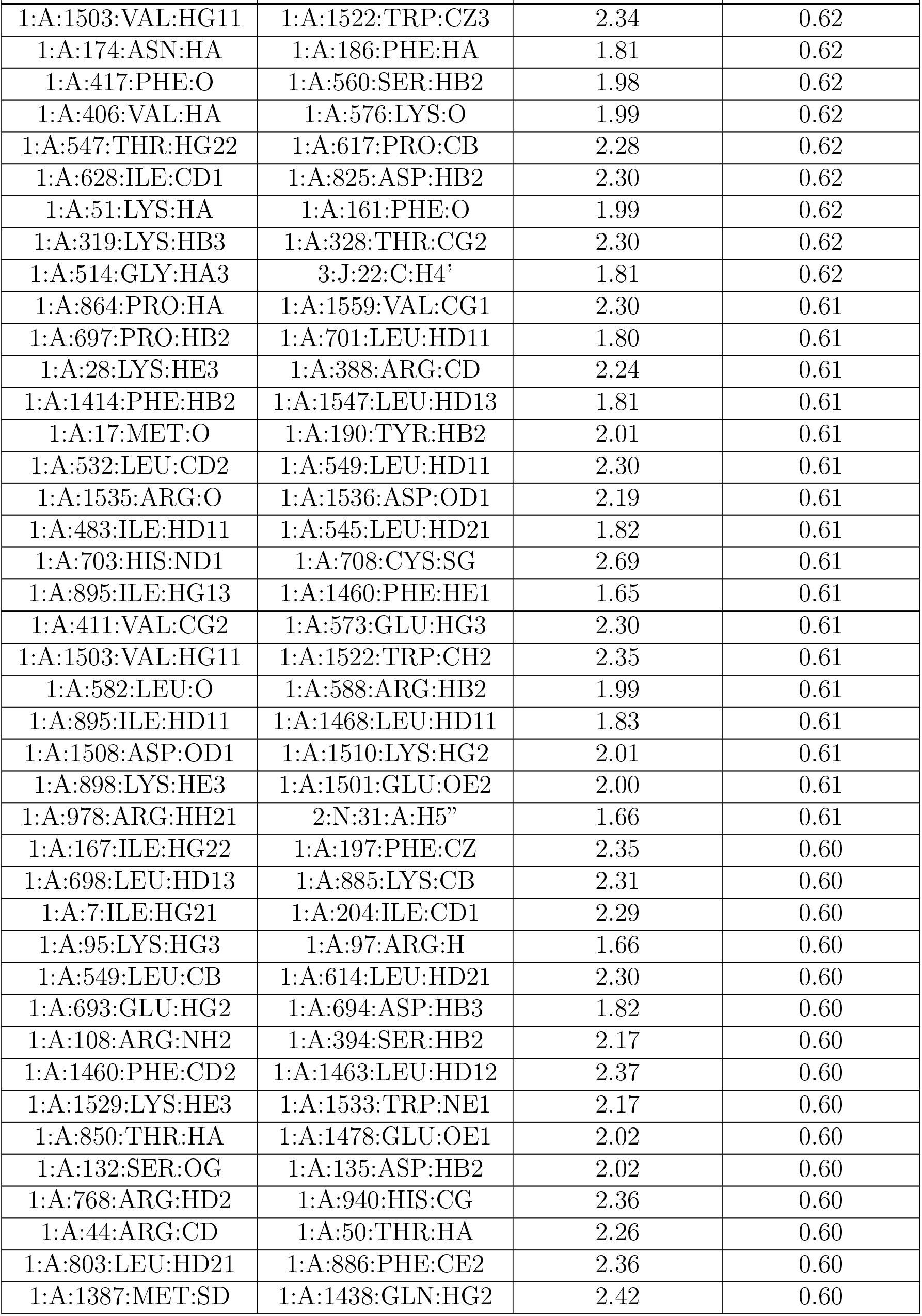
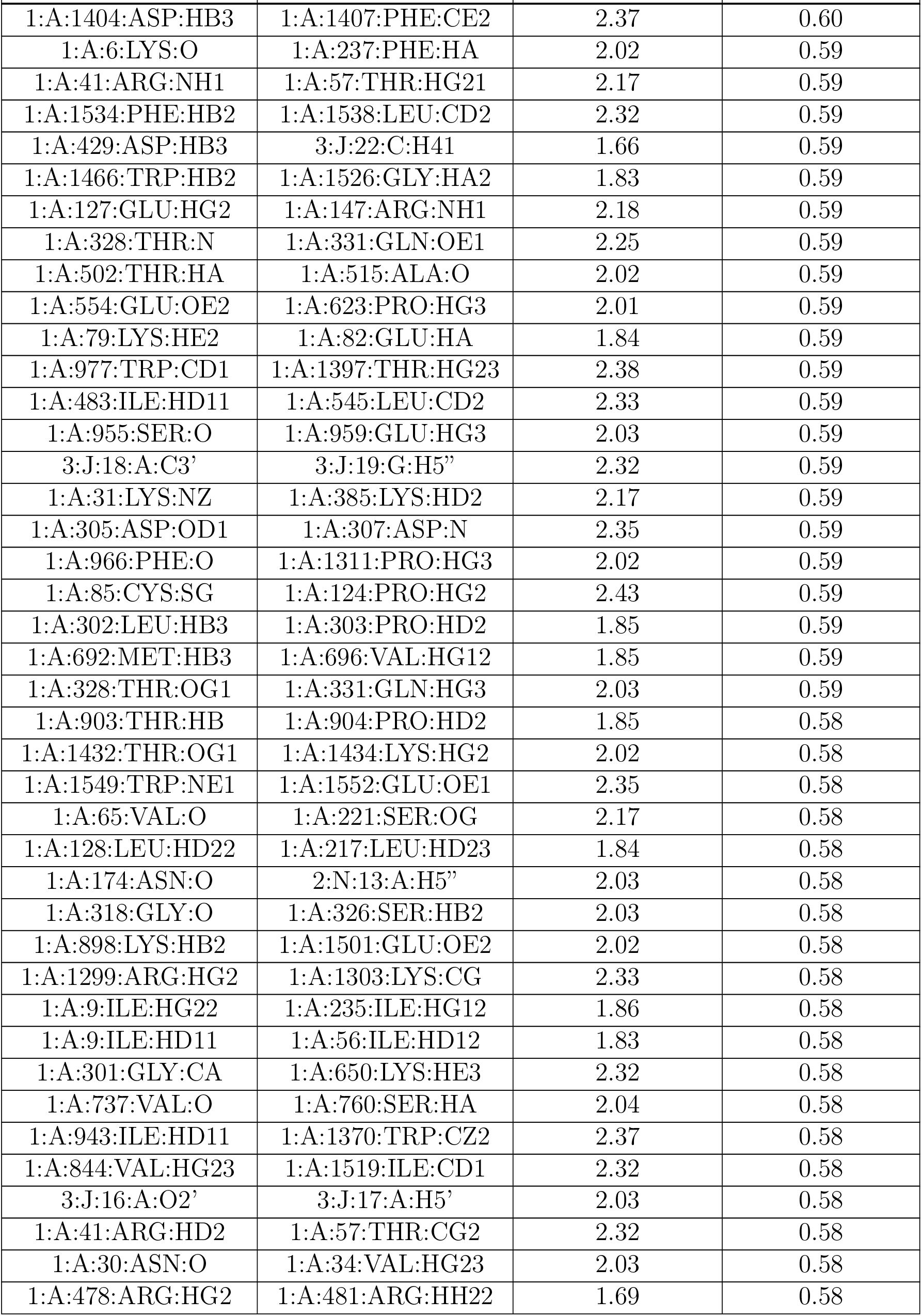
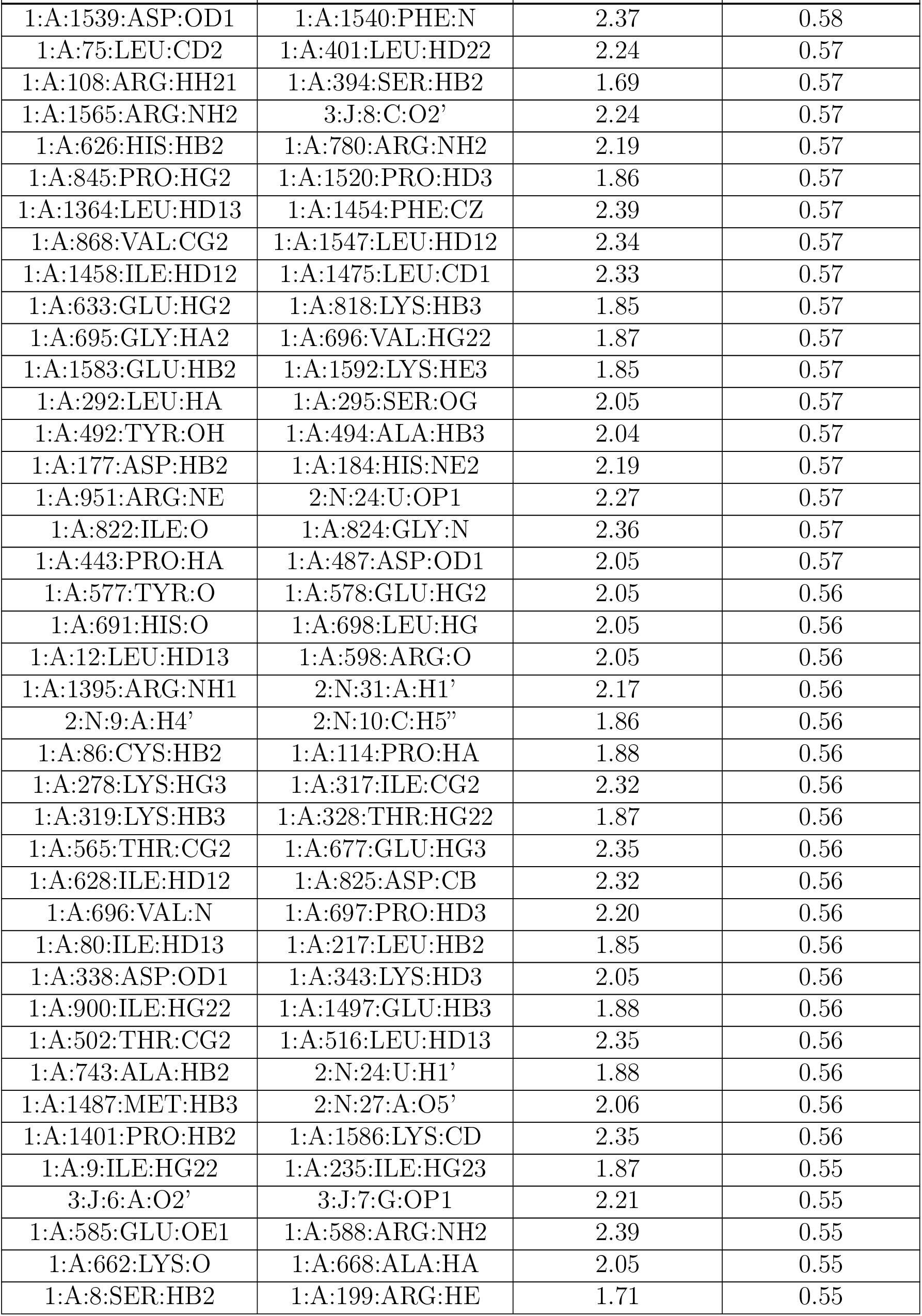
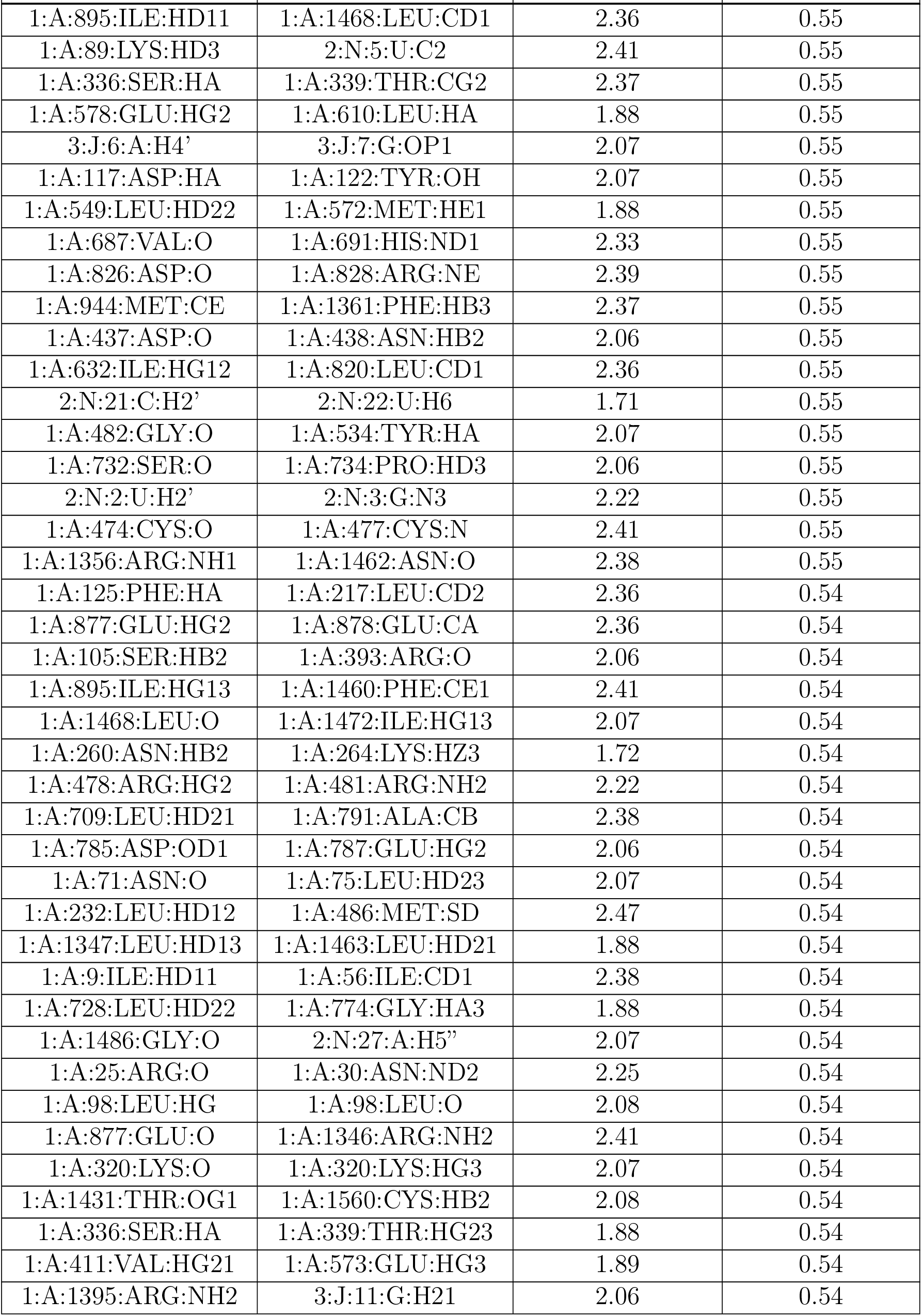
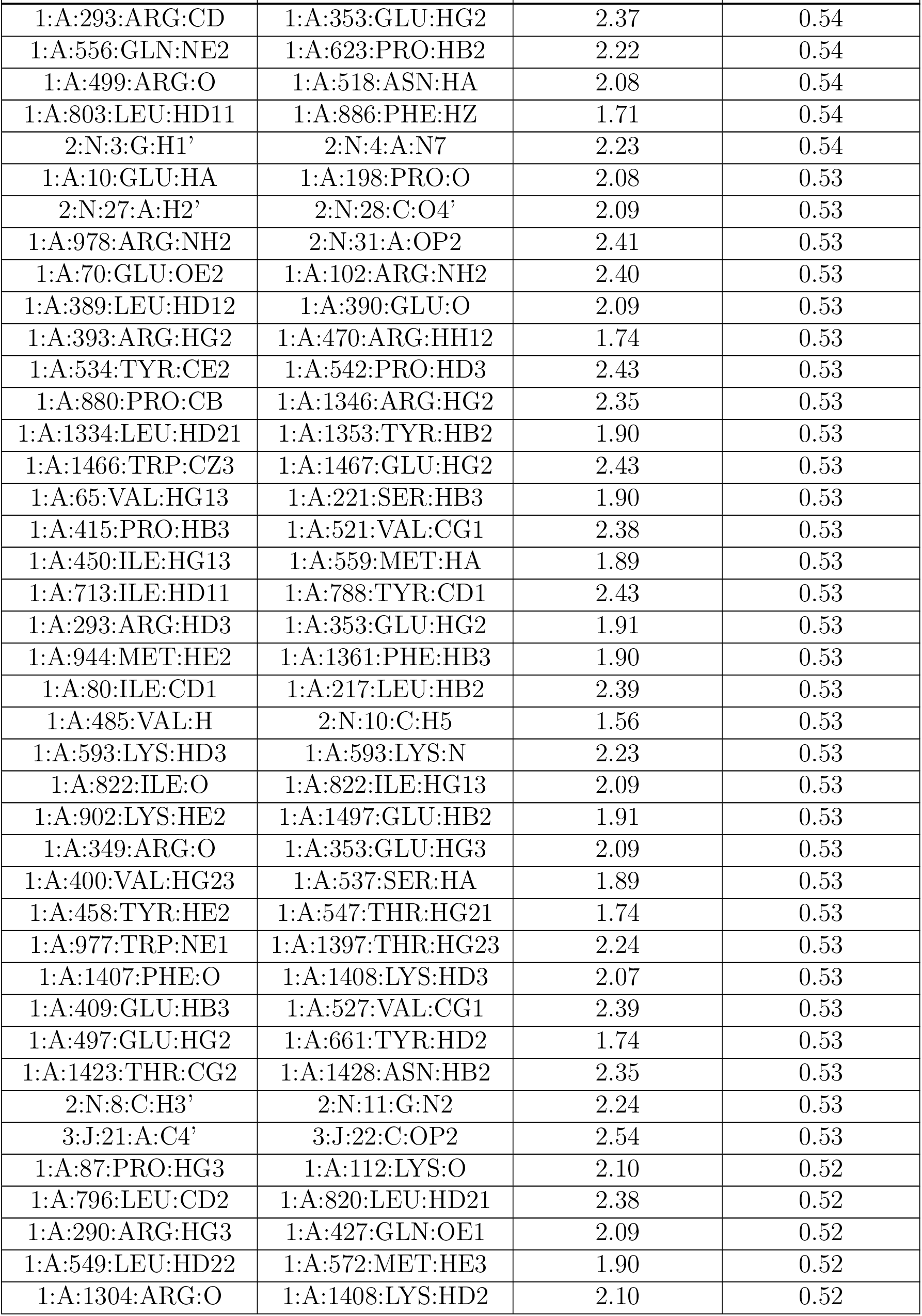
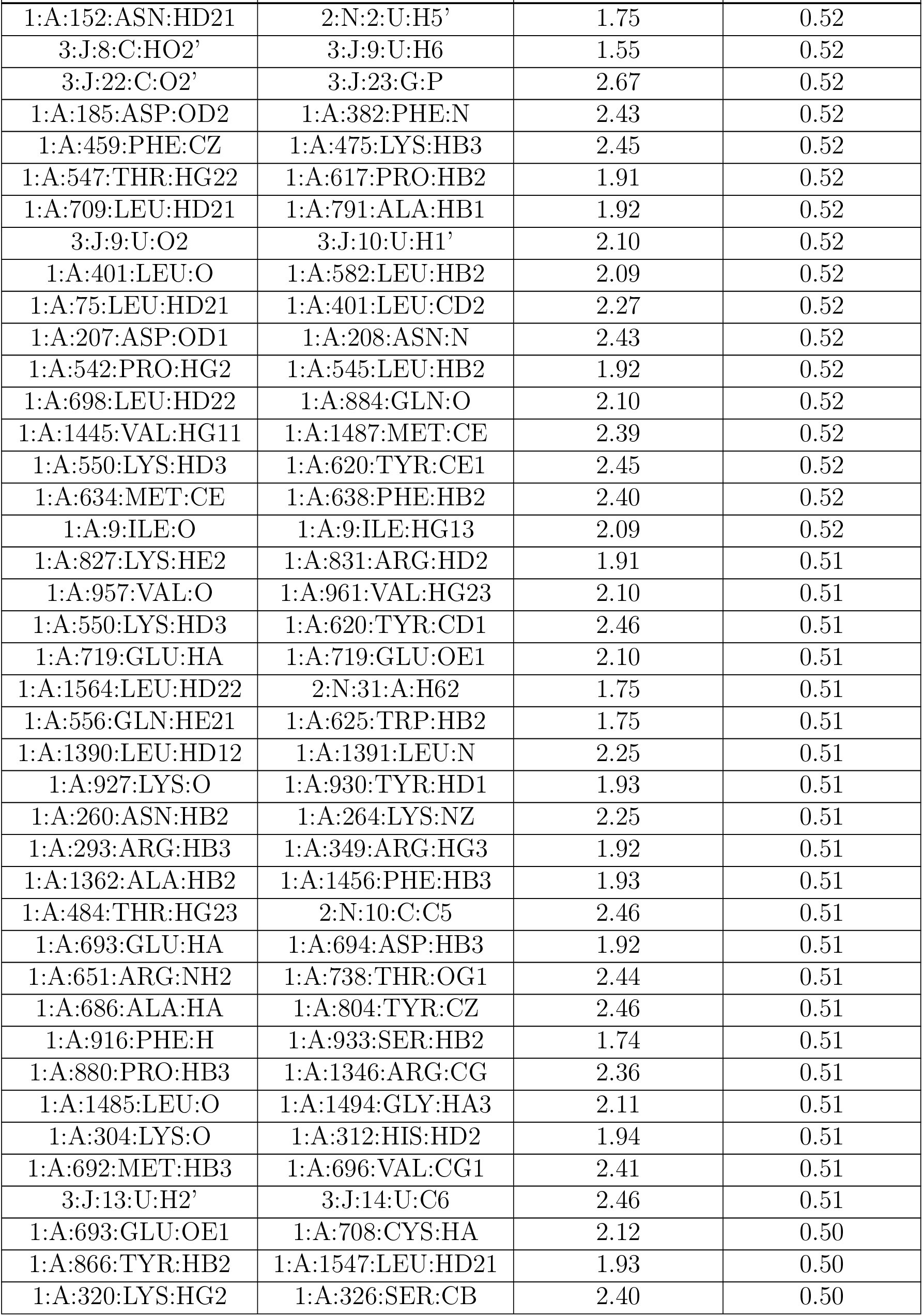
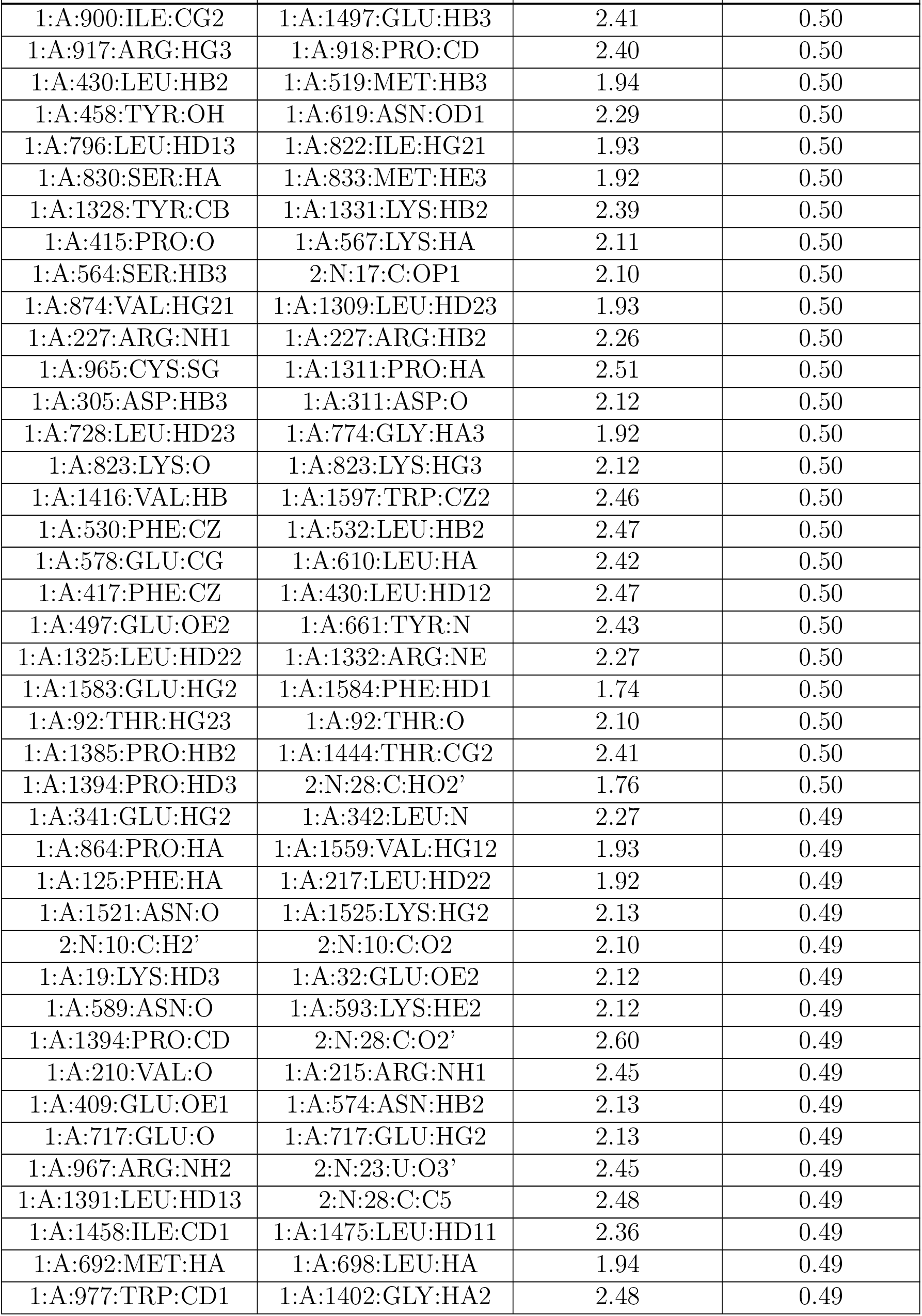
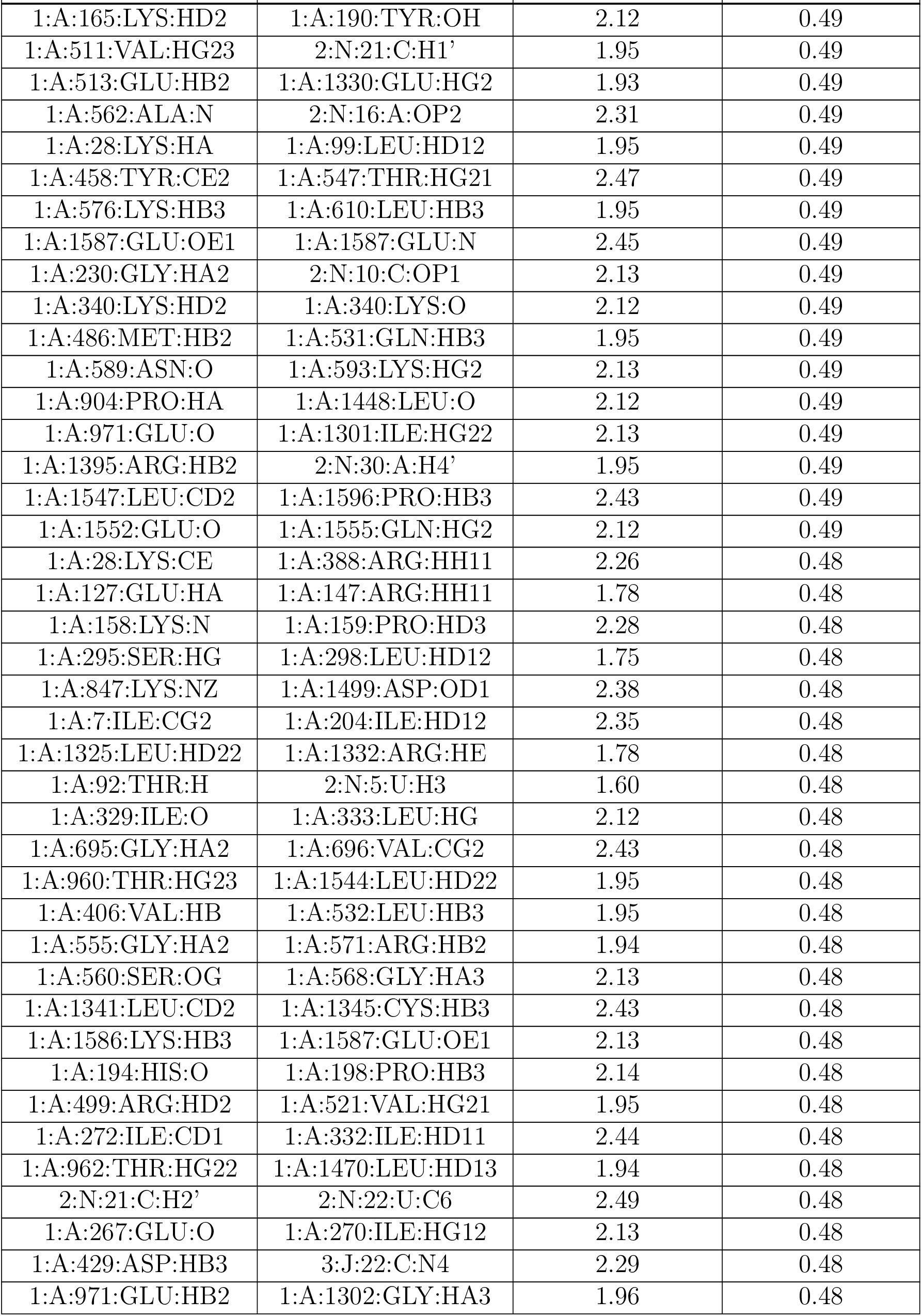
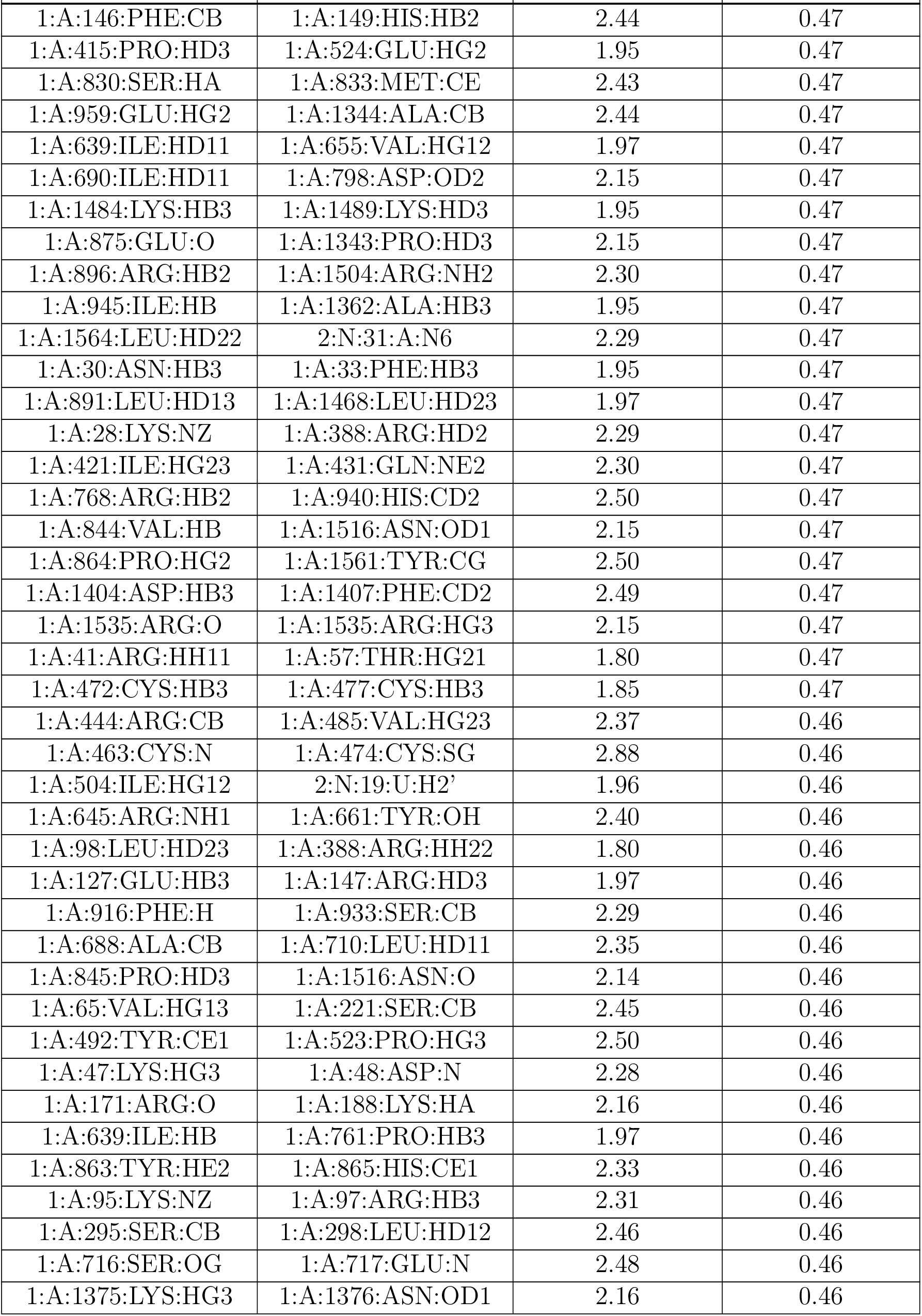
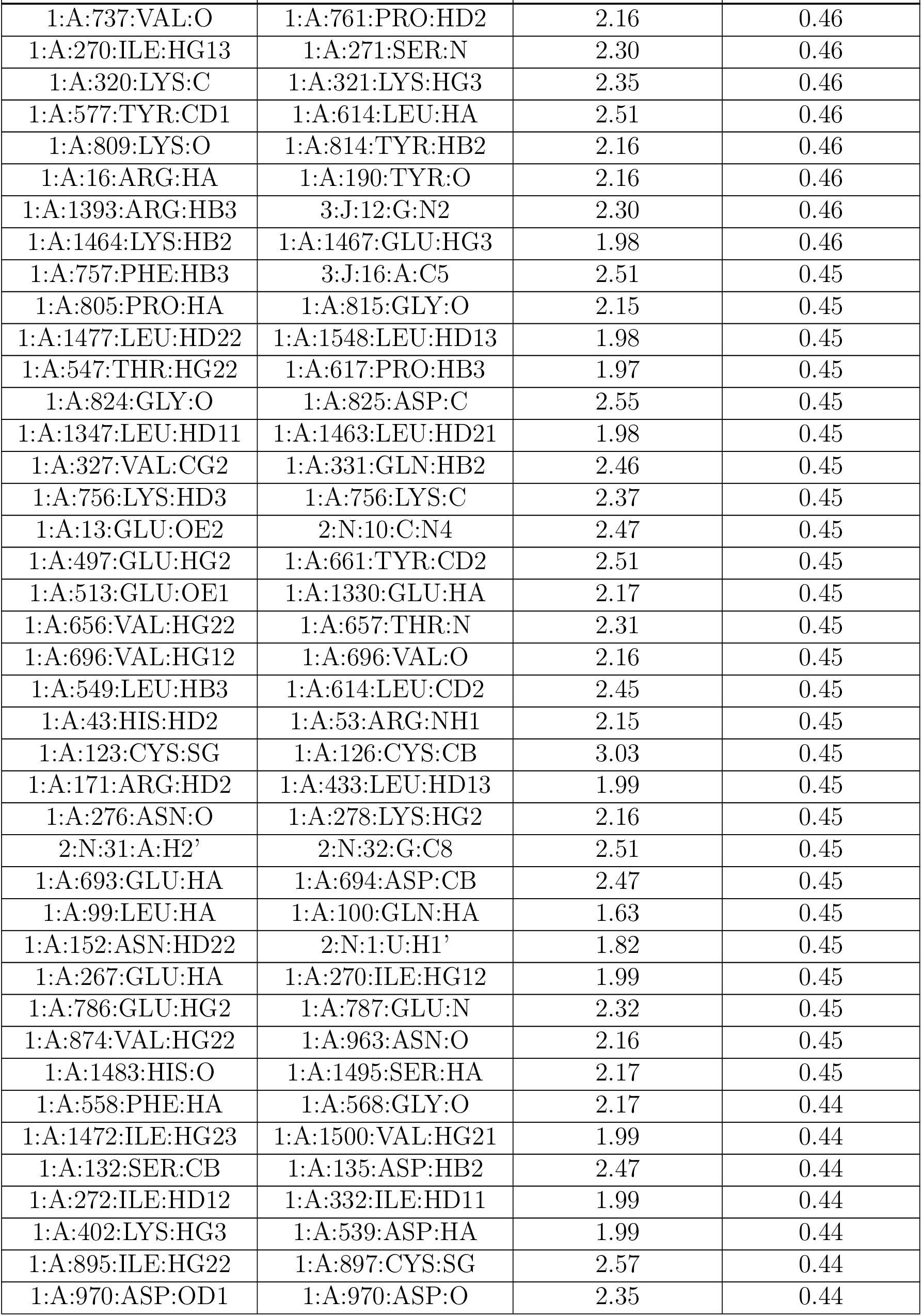
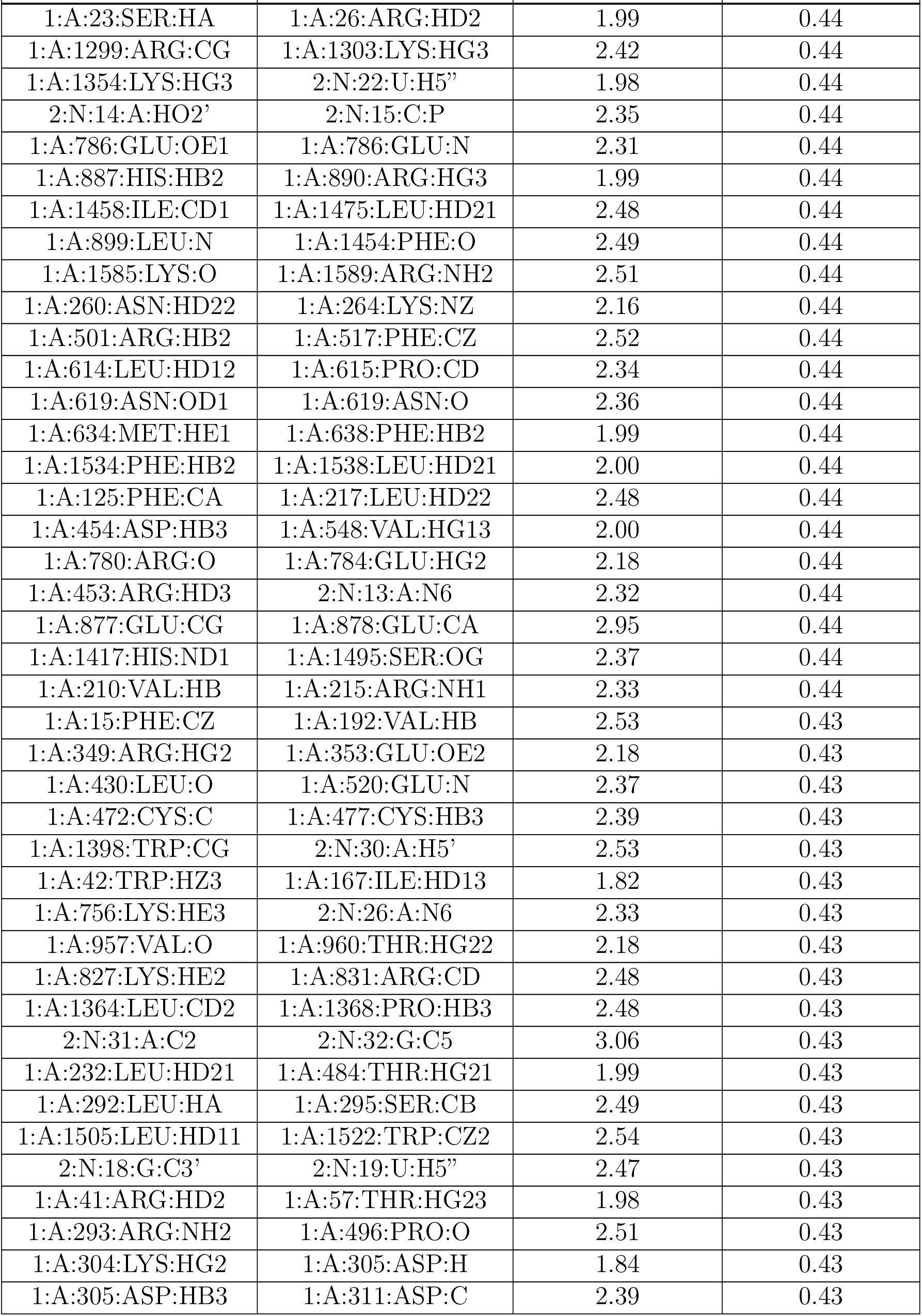
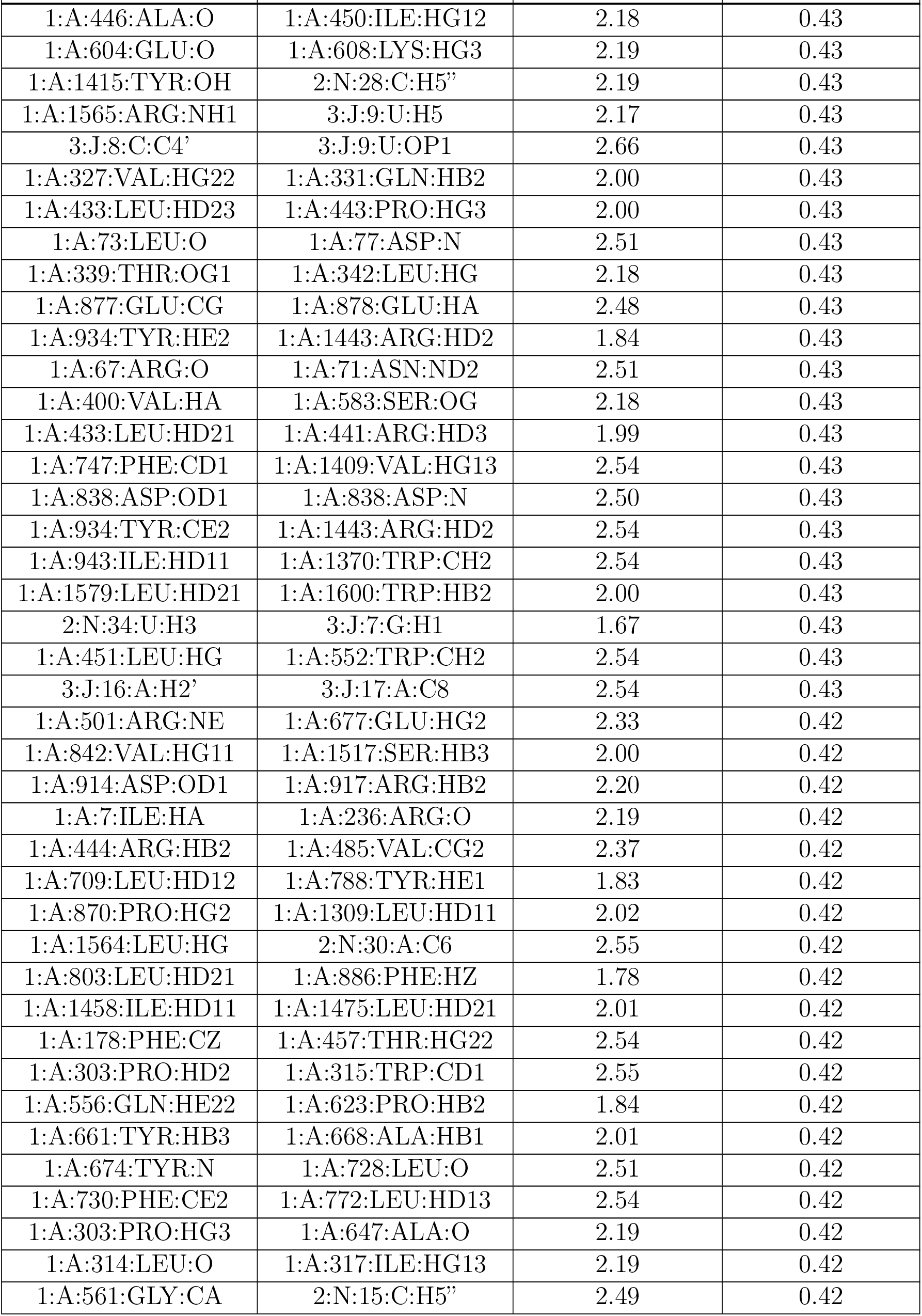
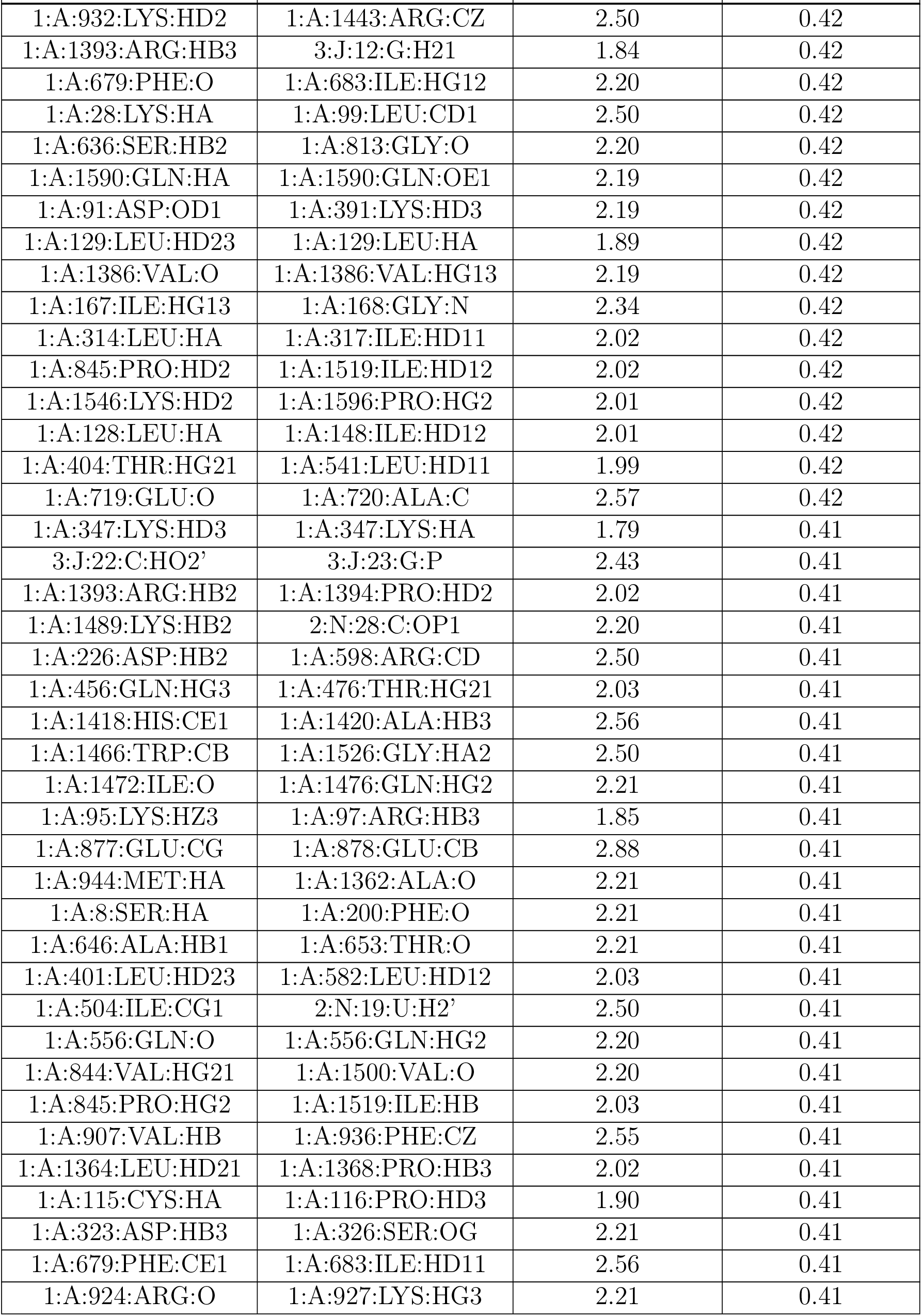
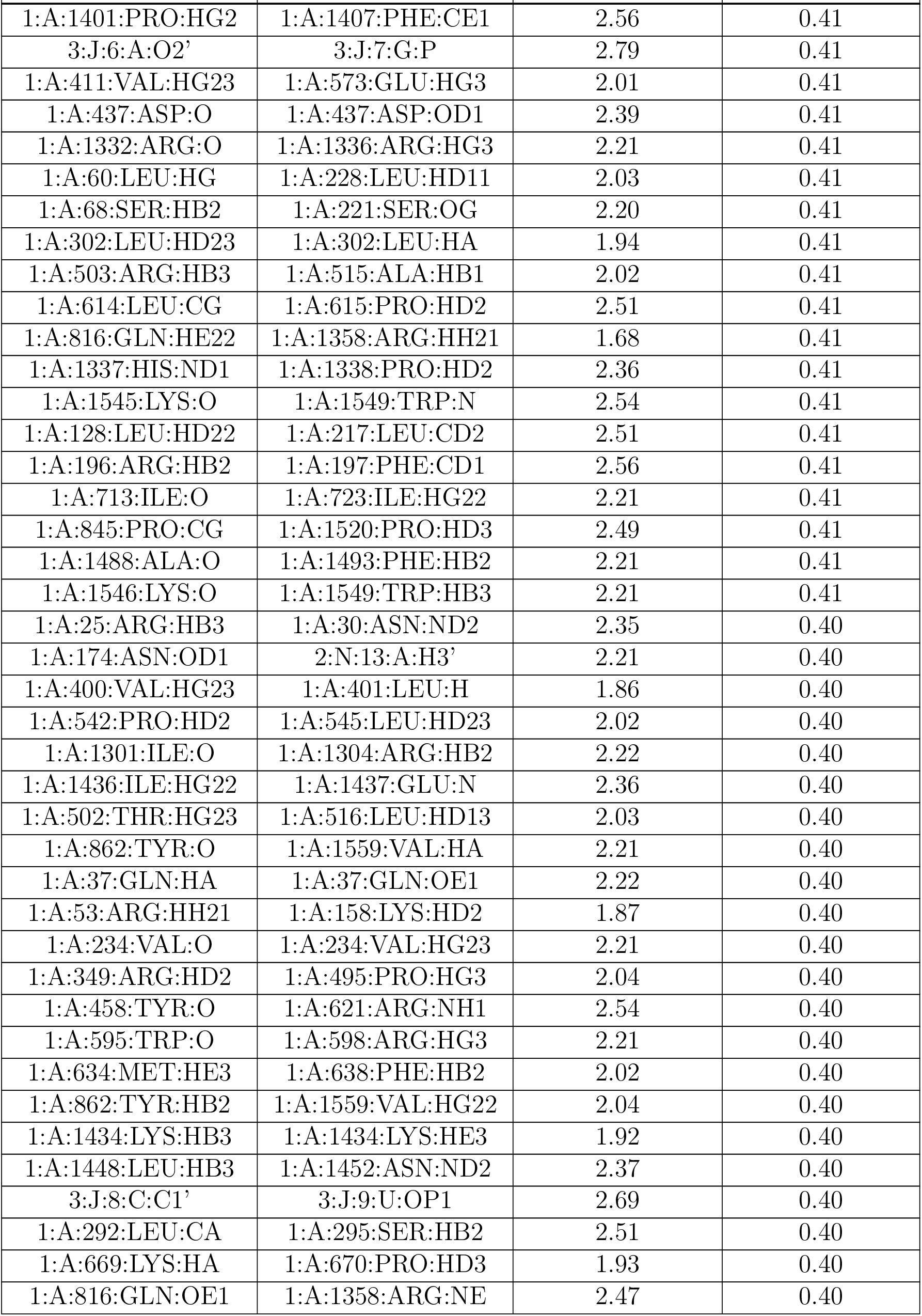
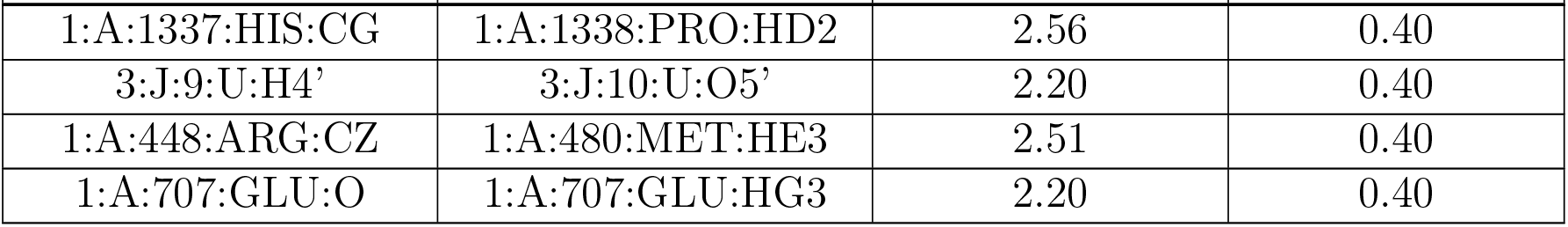

There are no symmetry-related clashes.

#### 5.3 Torsion angles

##### 5.3.1 Protein backbone

In the following table, the Percentiles column shows the percent Ramachandran outliers of the chain as a percentile score with respect to all PDB entries followed by that with respect to all EM entries.

The Analysed column shows the number of residues for which the backbone conformation was analysed, and the total number of residues.

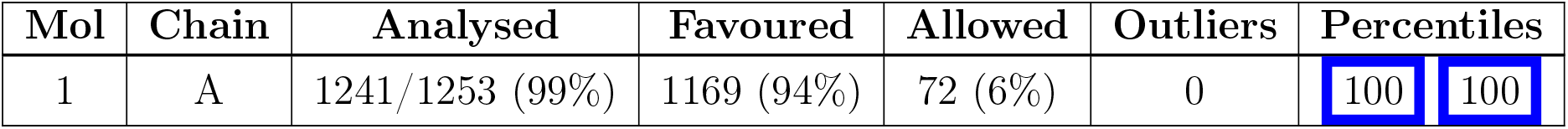

There are no Ramachandran outliers to report.

##### 5.3.2 Protein sidechains

In the following table, the Percentiles column shows the percent sidechain outliers of the chain as a percentile score with respect to all PDB entries followed by that with respect to all EM entries.

The Analysed column shows the number of residues for which the sidechain conformation was analysed, and the total number of residues.

| Mol | Chain | Analysed | Rotameric | Outliers | Percentiles |  |
| --- | --- | --- | --- | --- | --- | --- |
| 1 | A | 1075/1079 (100%) | 1073 (100%) | 2 (0%) | 93 | 98 |

All (2) residues with a non-rotameric sidechain are listed below:

| Mol | Chain | Res | Type |
| --- | --- | --- | --- |
| 1 | A | 321 | LYS |
| 1 | A | 640 | ASN |

Sometimes sidechains can be flipped to improve hydrogen bonding and reduce clashes. All (2) such sidechains are listed below:

| Mol | Chain | Res | Type |
| --- | --- | --- | --- |
| 1 | A | 556 | GLN |
| 1 | A | 712 | GLN |

##### 5.3.3 RNA

5.3.3 RNA i
| Mol | Chain | Analysed | Backbone Outliers | Pucker Outliers |
| --- | --- | --- | --- | --- |
| 2 | N | 33/34 (97%) | 12 (36%) | 0 |
| 3 | J | 18/18 (100%) | 9 (50%) | 4 (22%) |
| All | All | 51/52 (98%) | 21 (41%) | 4 (7%) |

All (21) RNA backbone outliers are listed below:

| Mol | Chain | Res | Type |
| --- | --- | --- | --- |
| 2 | N | 2 | U |
| 2 | N | 4 | A |
| 2 | N | 11 | G |
| 2 | N | 12 | G |
| 2 | N | 13 | A |
| 2 | N | 14 | A |
| 2 | N | 18 | G |
| 2 | N | 19 | U |
| 2 | N | 24 | U |
| 2 | N | 30 | A |
| 2 | N | 31 | A |
| 2 | N | 32 | G |
| 3 | J | 7 | G |
| 3 | J | 8 | C |
| 3 | J | 9 | U |
| 3 | J | 10 | U |
| 3 | J | 12 | G |
| 3 | J | 16 | A |
| 3 | J | 19 | G |
| 3 | J | 22 | C |
| 3 | J | 23 | G |

All (4) RNA pucker outliers are listed below:

| Mol | Chain | Res | Type |
| --- | --- | --- | --- |
| 3 | J | 6 | A |

| Mol | Chain | Res | Type |
| --- | --- | --- | --- |
| 3 | J | 8 | C |
| 3 | J | 9 | U |
| 3 | J | 21 | A |

#### 5.4 Non-standard residues in protein, DNA, RNA chains

There are no non-standard protein/DNA/RNA residues in this entry.

#### 5.5 Carbohydrates

There are no monosaccharides in this entry.

#### 5.6 Ligand geometry

There are no ligands in this entry.

#### 5.7 Other polymers

There are no such residues in this entry.

#### 5.8 Polymer linkage issues

The following chains have linkage breaks:

| Mol | Chain | Number of breaks |
| --- | --- | --- |
| 1 | A | 3 |

All chain breaks are listed below:

| Model | Chain | Residue-1 | Atom-1 | Residue-2 | Atom-2 | Distance (Å) |
| --- | --- | --- | --- | --- | --- | --- |
| 1 | A | 238:ASP | C | 257:PRO | N | 44.03 |
| 1 | A | 141:LYS | C | 142:LYS | N | 10.82 |
| 1 | A | 978:ARG | C | 1296:VAL | N | 5.59 |

### 6 Map visualisation

This section contains visualisations of the EMDB entry EMD-27138. These allow visual inspection of the internal detail of the map and identification of artifacts.

Images derived from a raw map, generated by summing the deposited half-maps, are presented below the corresponding image components of the primary map to allow further visual inspection and comparison with those of the primary map.

#### 6.1 Orthogonal projections

##### 6.1.1 Primary map

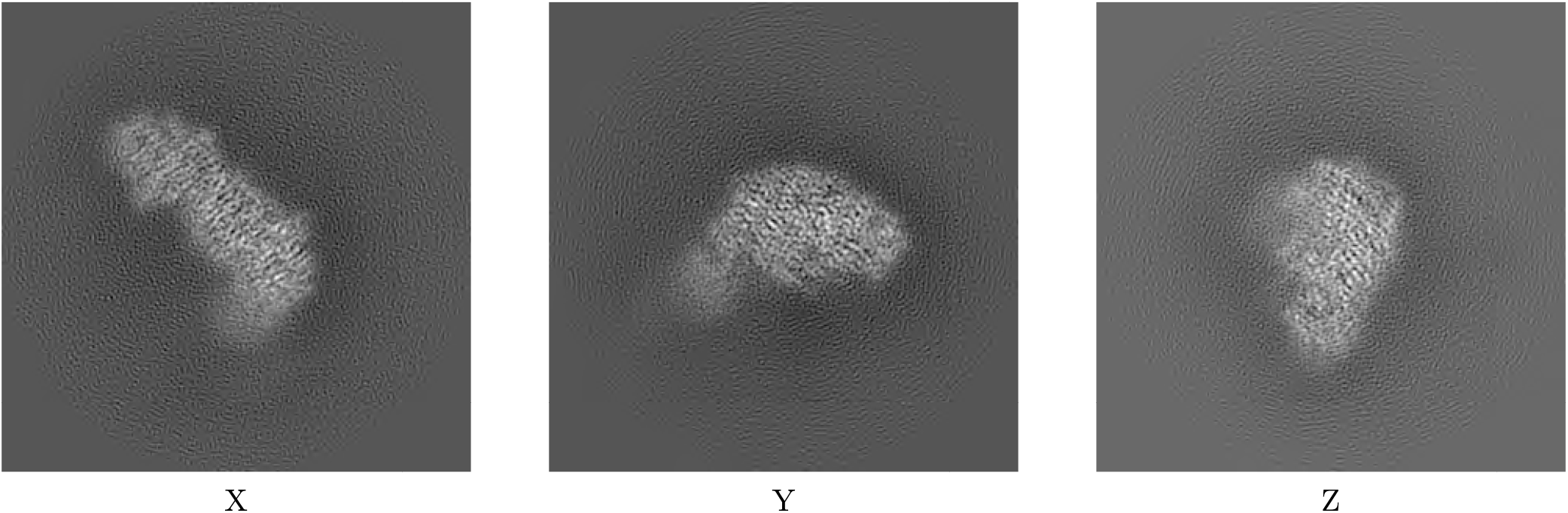

##### 6.1.2 Raw map

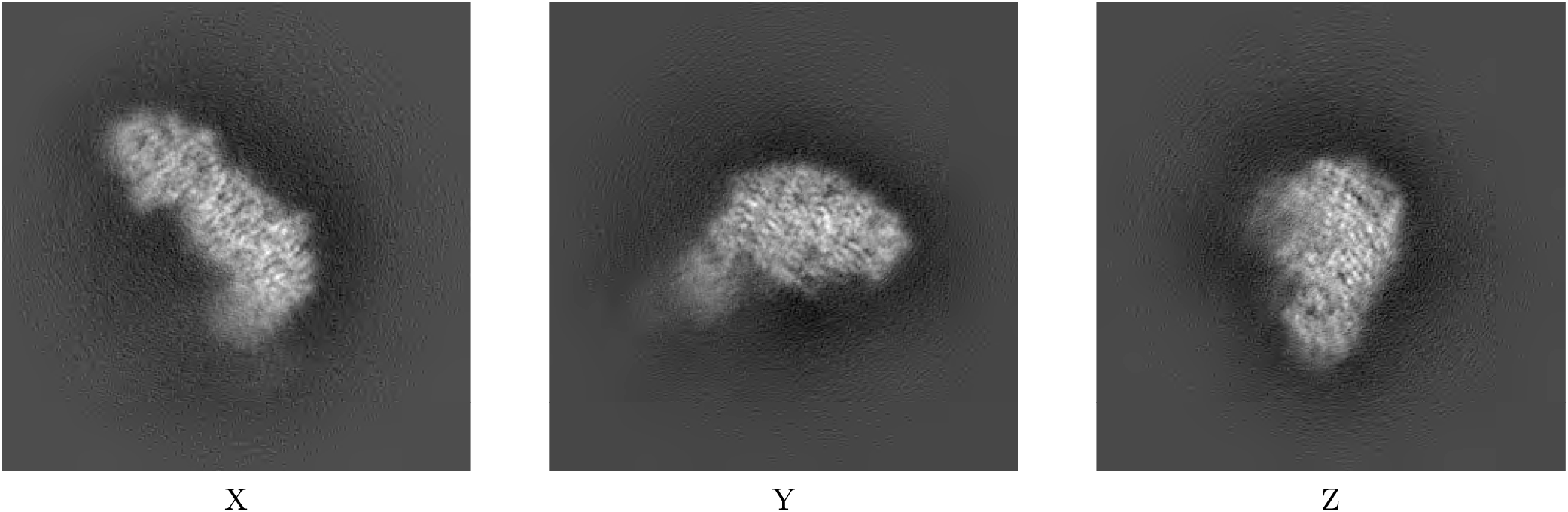

The images above show the map projected in three orthogonal directions.

#### 6.2 Central slices

##### 6.2.1 Primary map

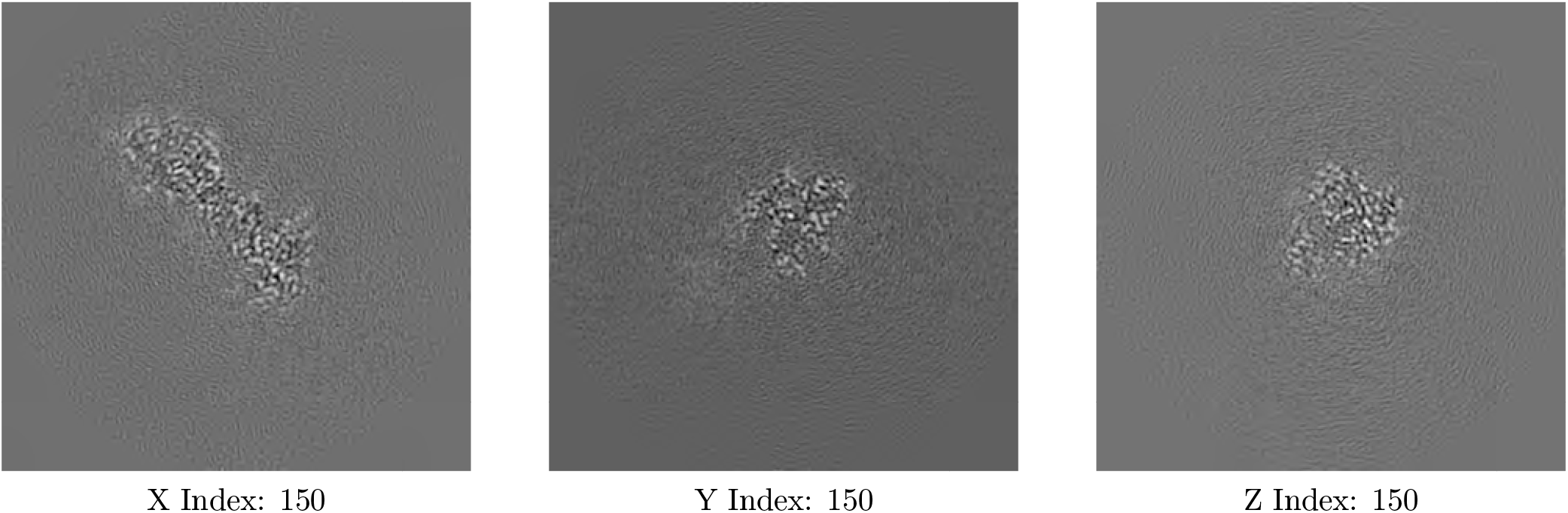

##### 6.2.2 Raw map

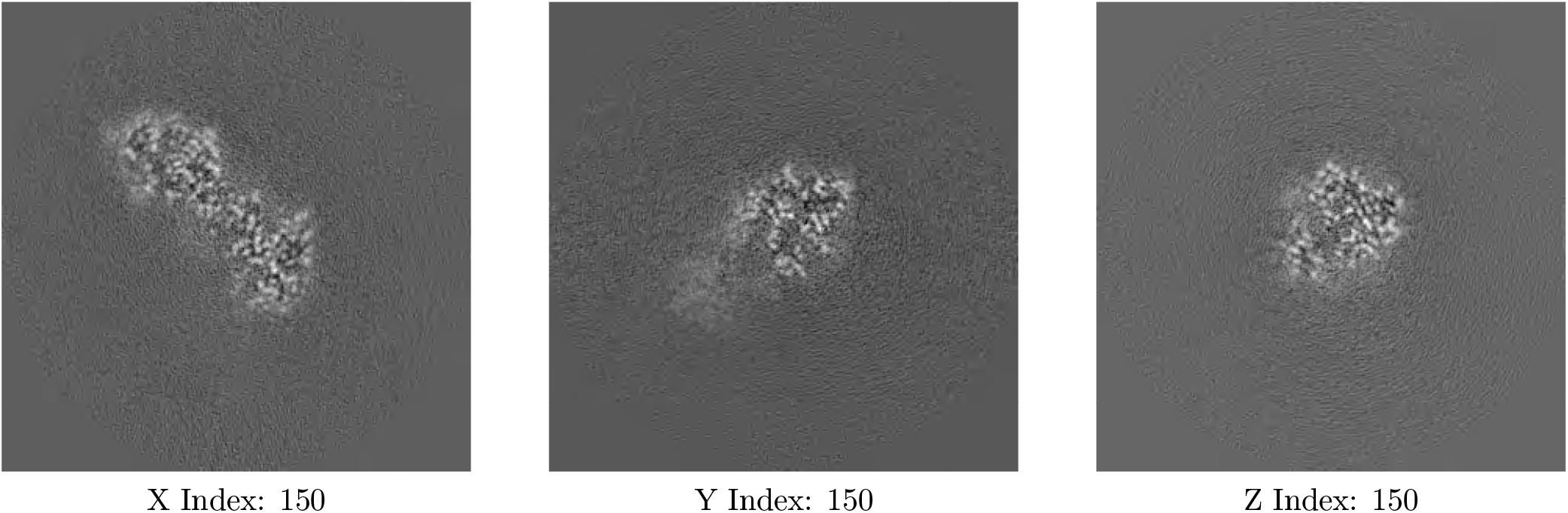

The images above show central slices of the map in three orthogonal directions.

#### 6.3 Largest variance slices

##### 6.3.1 Primary map

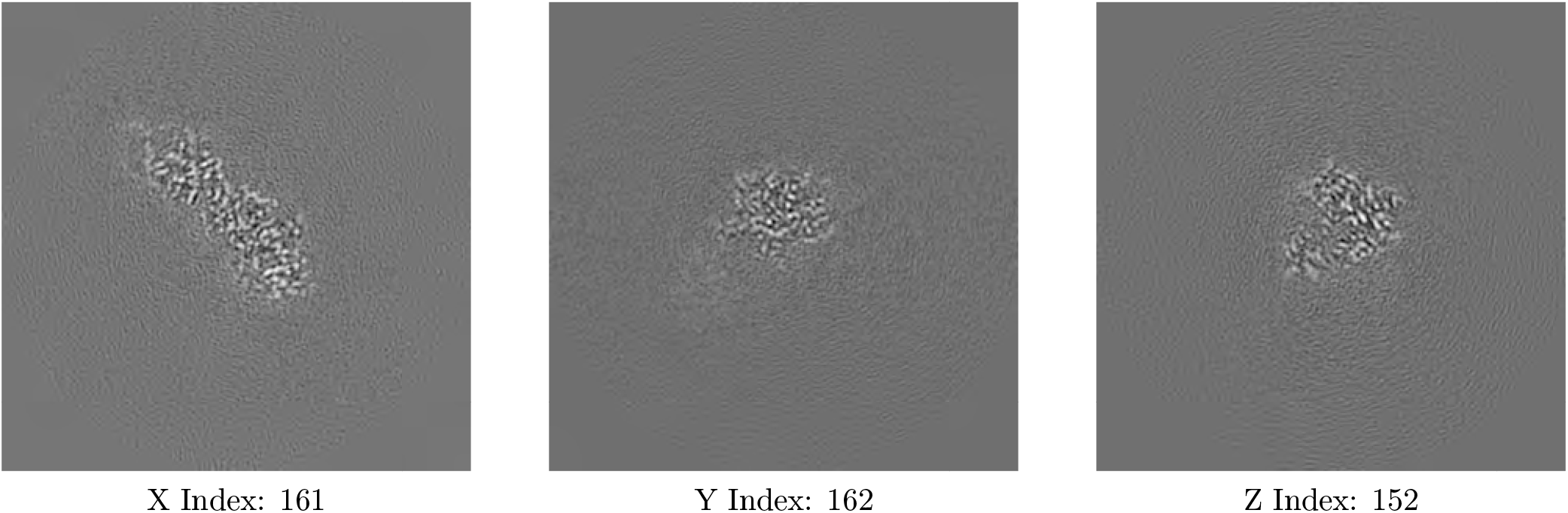

##### 6.3.2 Raw map

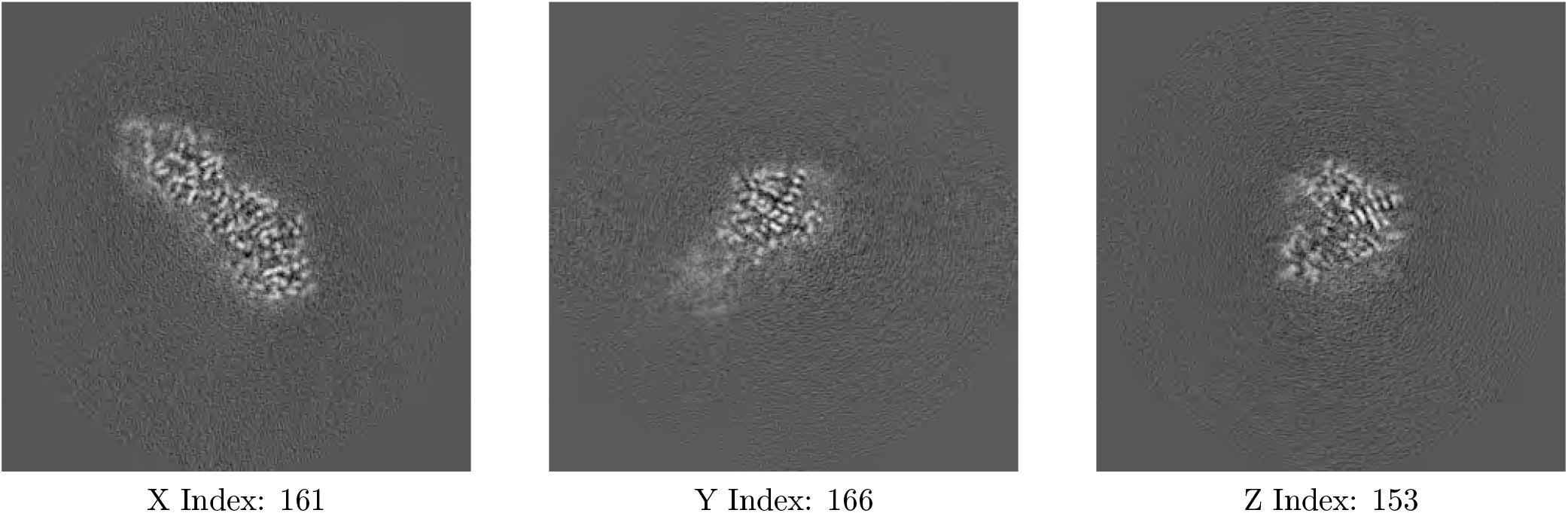

The images above show the largest variance slices of the map in three orthogonal directions.

#### 6.4 Orthogonal surface views

##### 6.4.1 Primary map

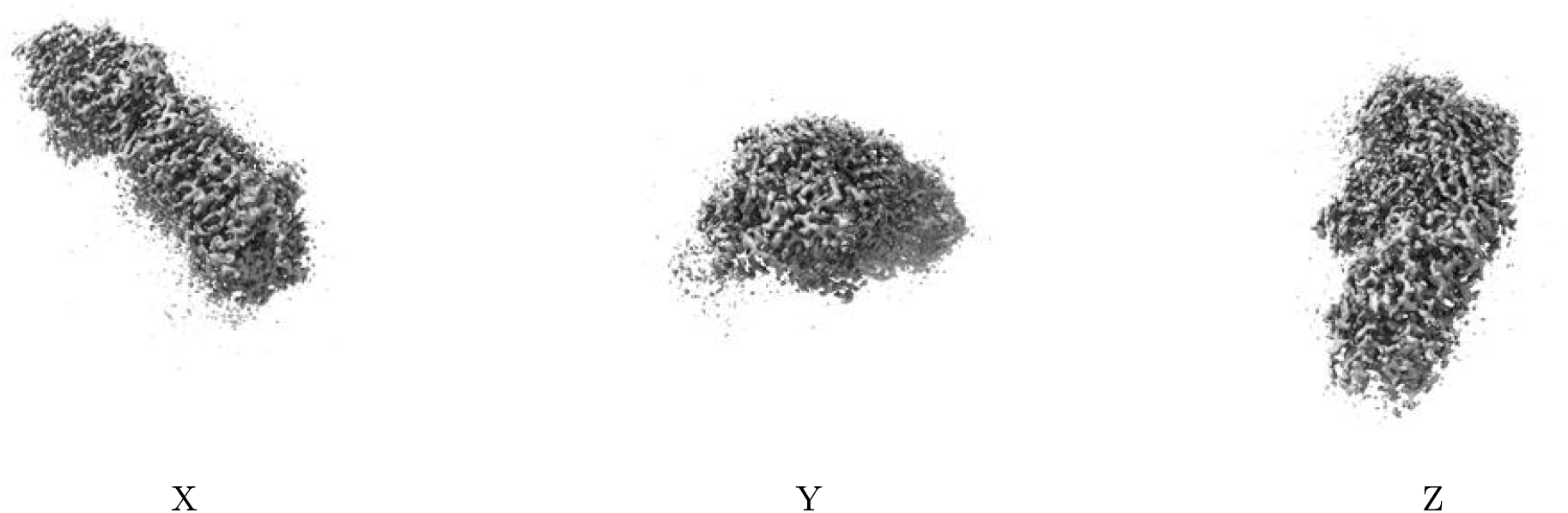

The images above show the 3D surface view of the map at the recommended contour level 0.0146. These images, in conjunction with the slice images, may facilitate assessment of whether an appropriate contour level has been provided.

##### 6.4.2 Raw map

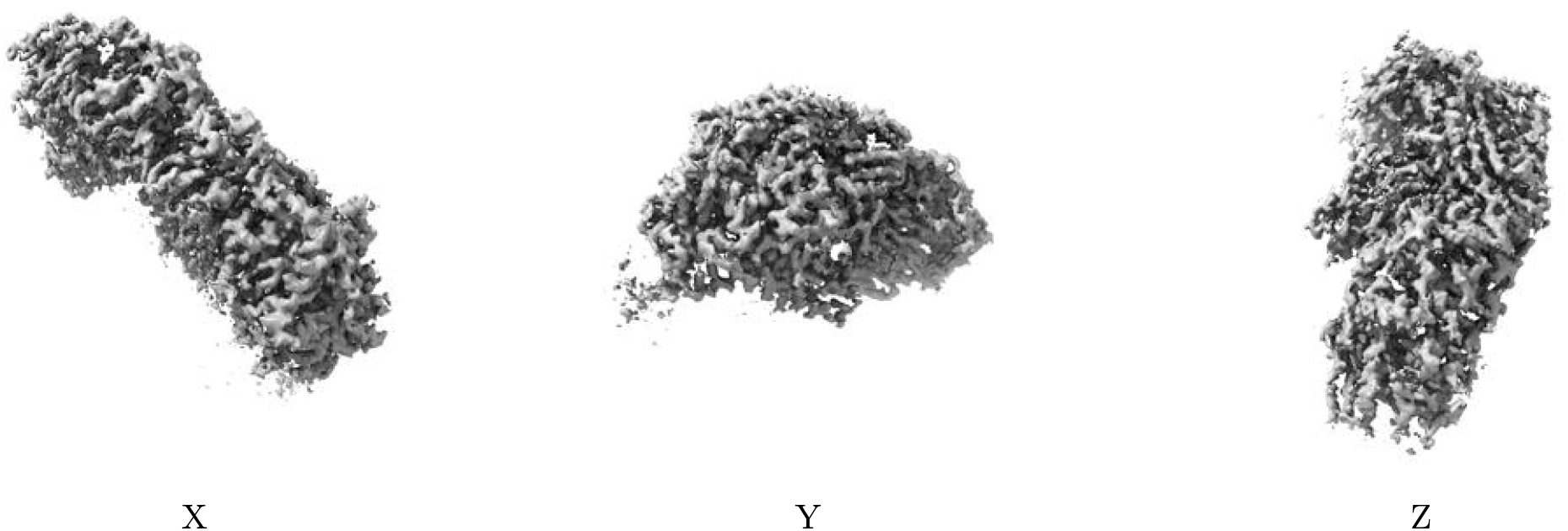

These images show the 3D surface of the raw map. The raw map’s contour level was selected so that its surface encloses the same volume as the primary map does at its recommended contour level.

#### 6.5 Mask visualisation

### 7 Map analysis

This section contains the results of statistical analysis of the map.

#### 7.1 Map-value distribution

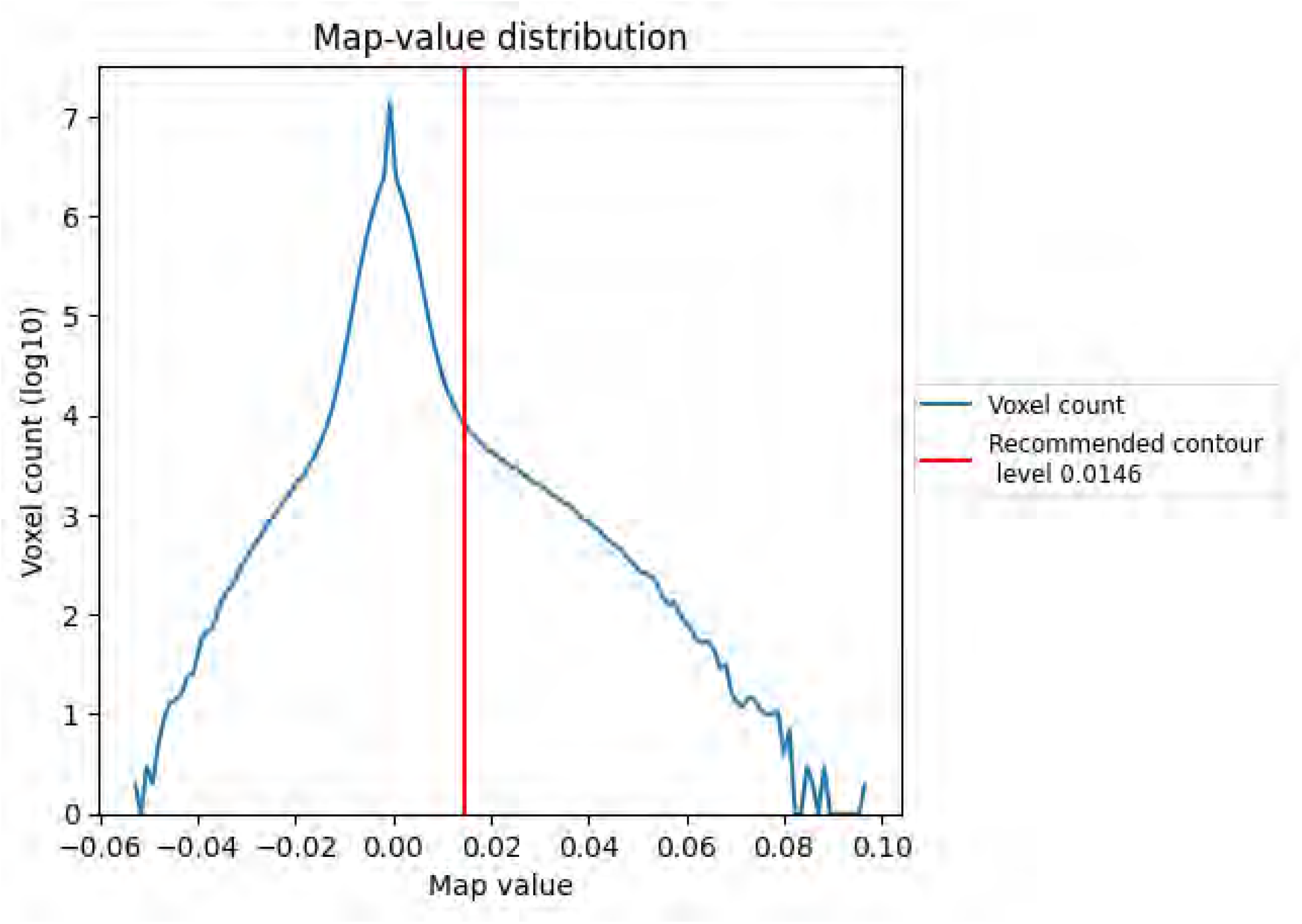

The map-value distribution is plotted in 128 intervals along the x-axis. The y-axis is logarithmic. A spike in this graph at zero usually indicates that the volume has been masked.

#### 7.2 Volume estimate

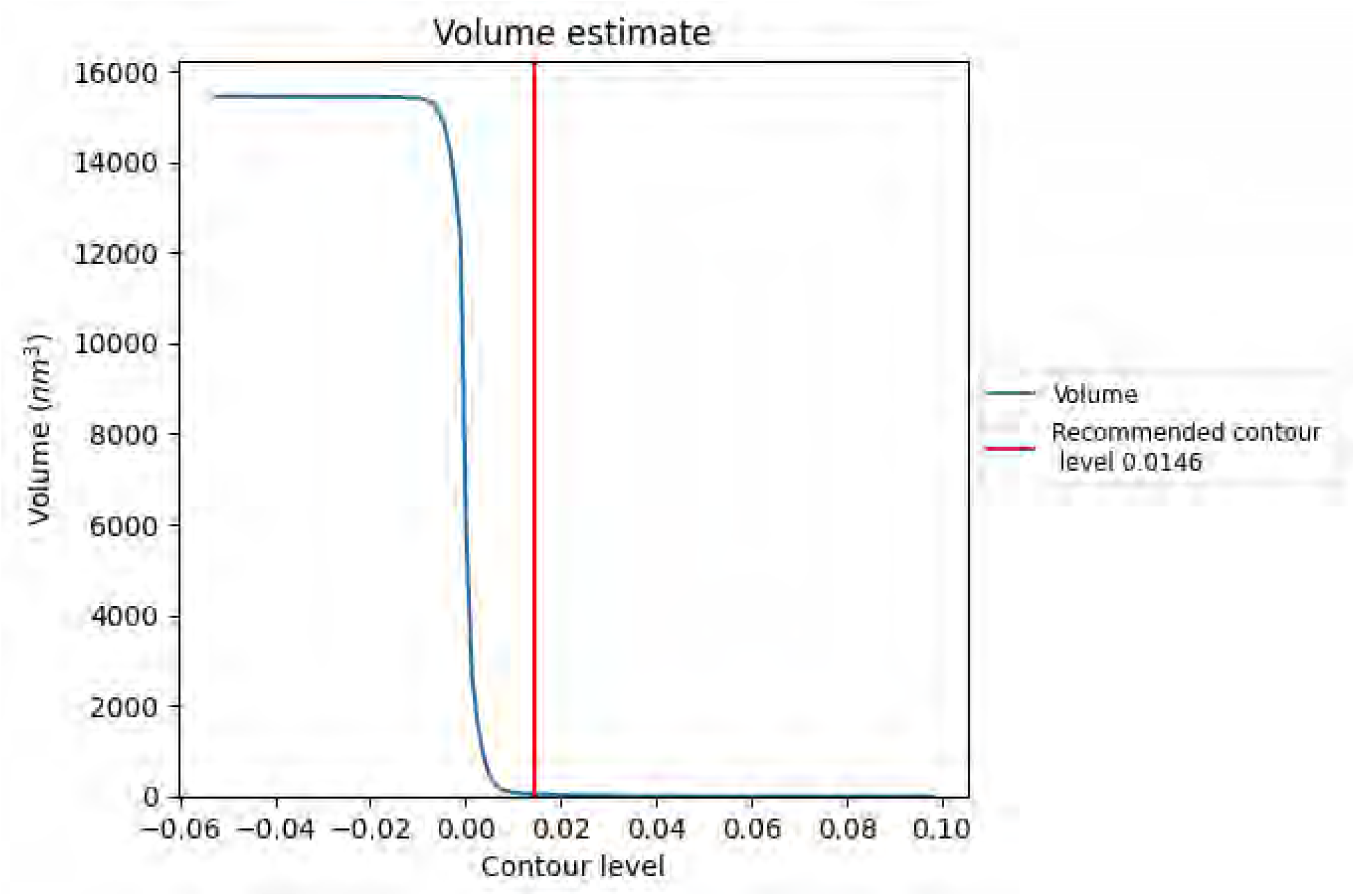

The volume at the recommended contour level is 43 nm^3^; this corresponds to an approximate mass of 39 kDa.

The volume estimate graph shows how the enclosed volume varies with the contour level. The recommended contour level is shown as a vertical line and the intersection between the line and the curve gives the volume of the enclosed surface at the given level.

#### 7.3 Rotationally averaged power spectrum

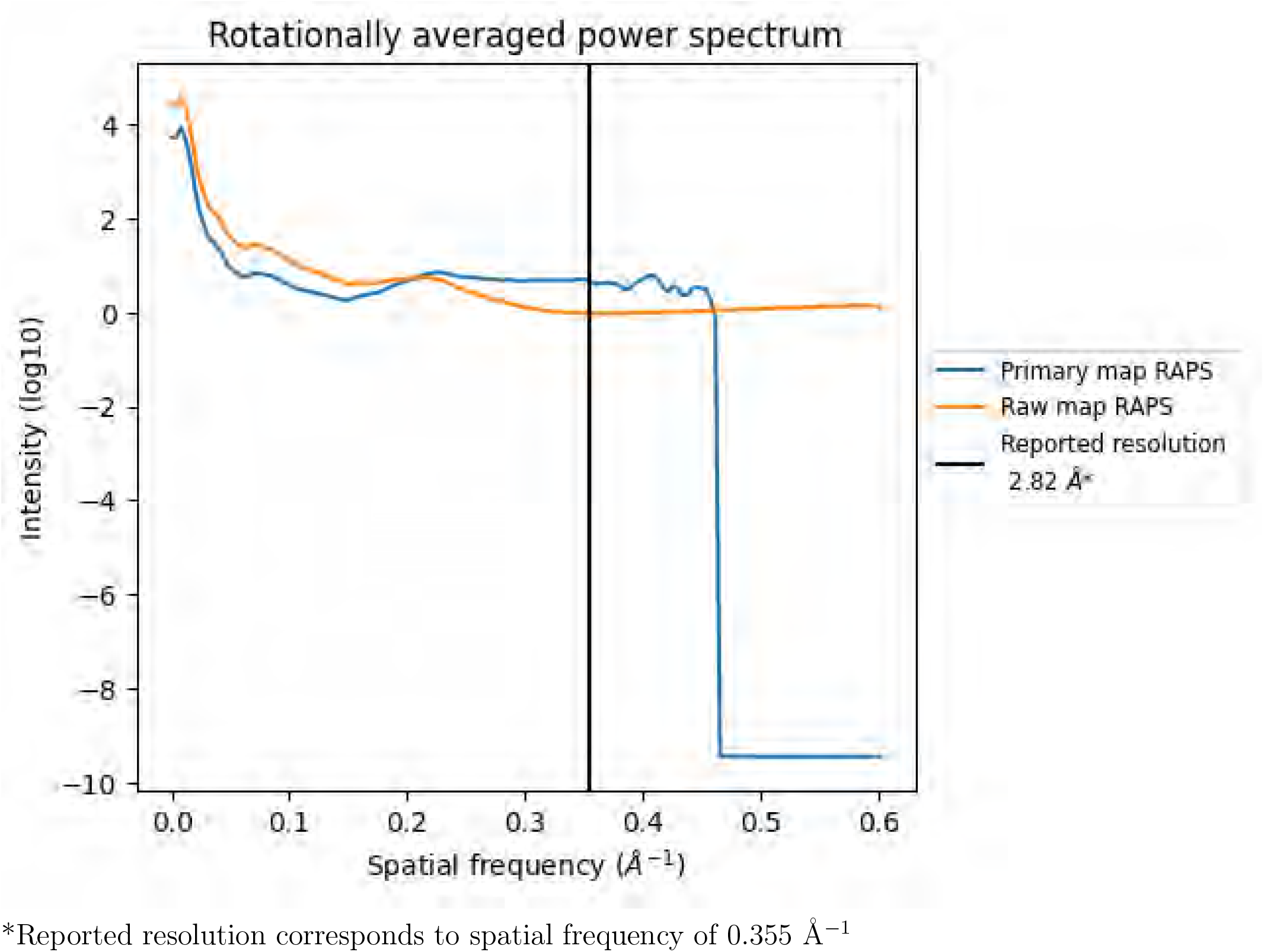

### 8 Fourier-Shell correlation

Fourier-Shell Correlation (FSC) is the most commonly used method to estimate the resolution of single-particle and subtomogram-averaged maps. The shape of the curve depends on the imposed symmetry, mask and whether or not the two 3D reconstructions used were processed from a common reference. The reported resolution is shown as a black line. A curve is displayed for the half-bit criterion in addition to lines showing the 0.143 gold standard cut-off and 0.5 cut-off.

#### 8.1 FSC

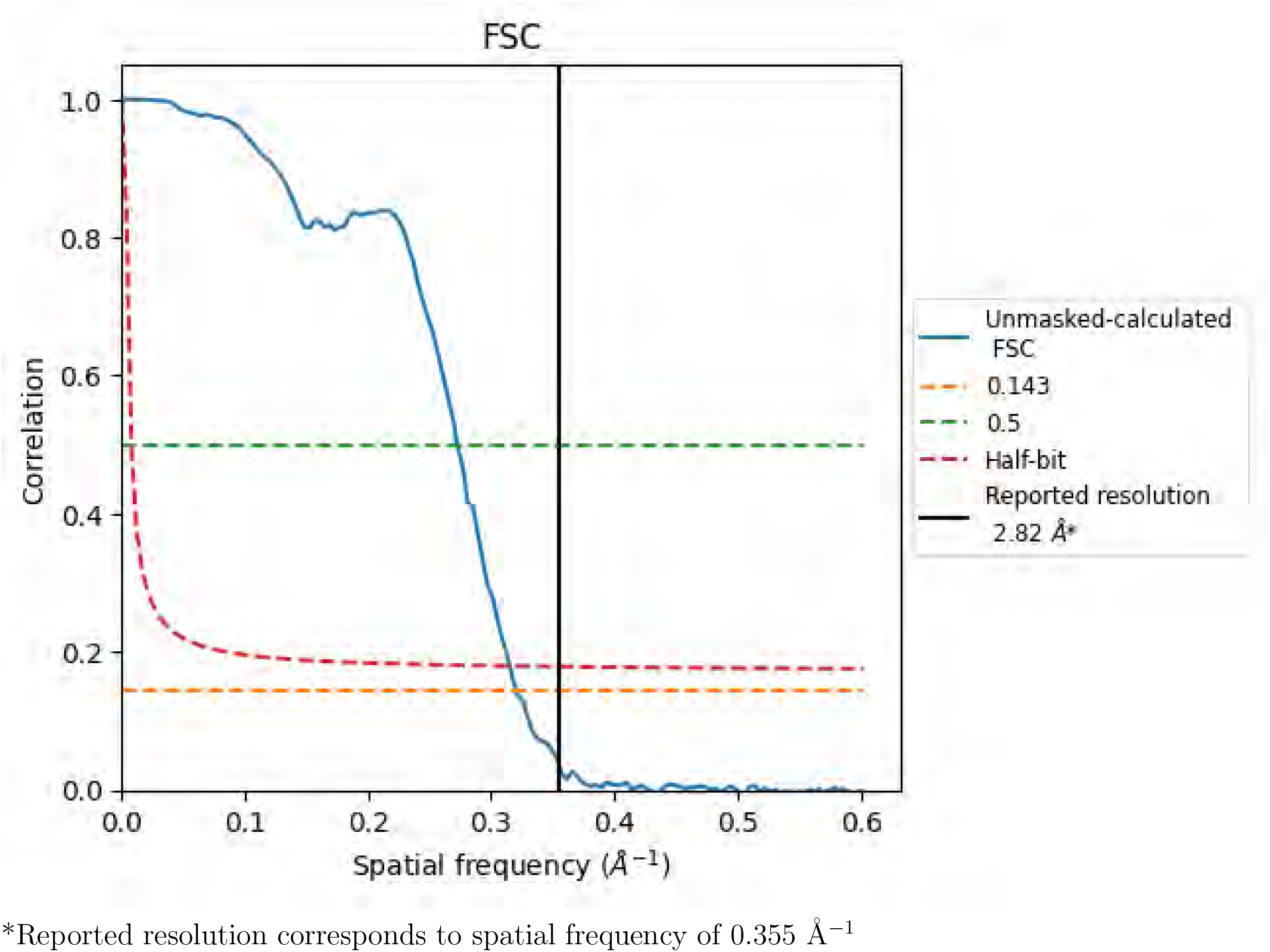

#### 8.2 Resolution estimates

8.2 Resolution estimates i
| Resolution estimate (Å) | Estimation criterion (FSC cut-off) |  |  |
| --- | --- | --- | --- |
|  | 0.143 | 0.5 | Half-bit |
| Reported by author | 2.82 | - | - |
| Author-provided FSC curve | - | - | - |
| Unmasked-calculated* | 3.12 | 3.67 | 3.17 |

\*Resolution estimate based on FSC curve calculated by comparison of deposited half-maps. The value from deposited half-maps intersecting FSC 0.143 CUT-OFF 3.12 differs from the reported value 2.82 by more than 10 %

### 9 Map-model fit

This section contains information regarding the fit between EMDB map EMD-27138 and PDB model 8D1V. Per-residue inclusion information can be found in section 3 on page 4.

#### 9.1 Map-model overlay

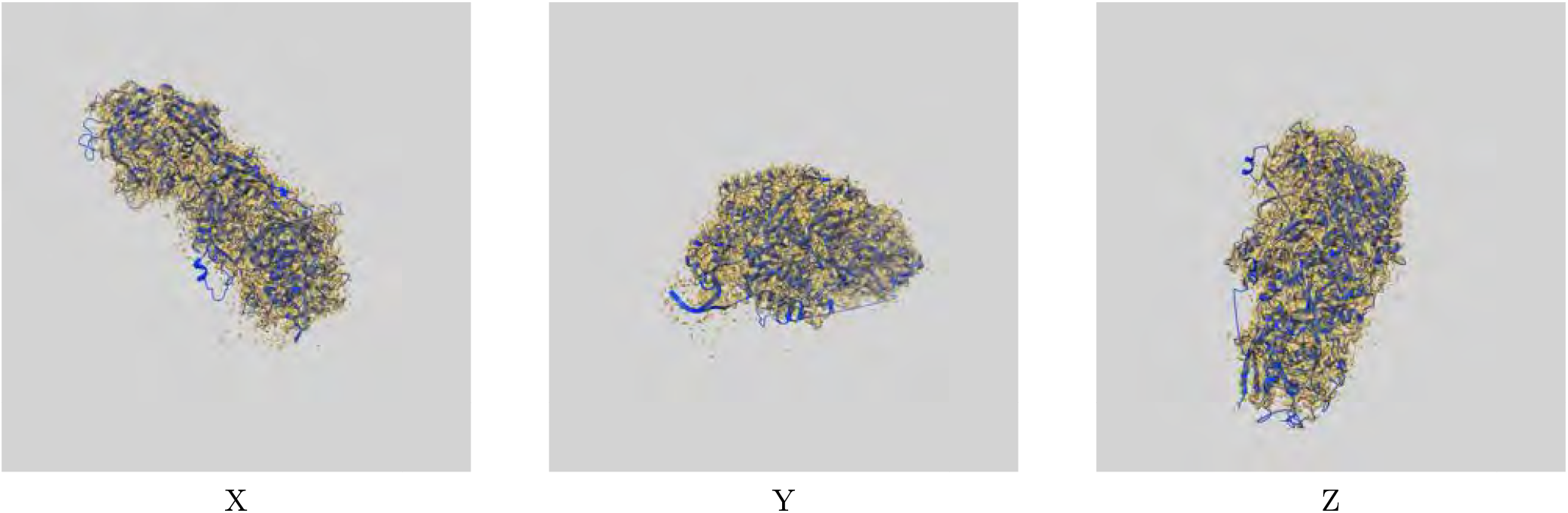

The images above show the 3D surface view of the map at the recommended contour level 0.0146 at 50% transparency in yellow overlaid with a ribbon representation of the model coloured in blue. These images allow for the visual assessment of the quality of fit between the atomic model and the map.

#### 9.2 Atom inclusion

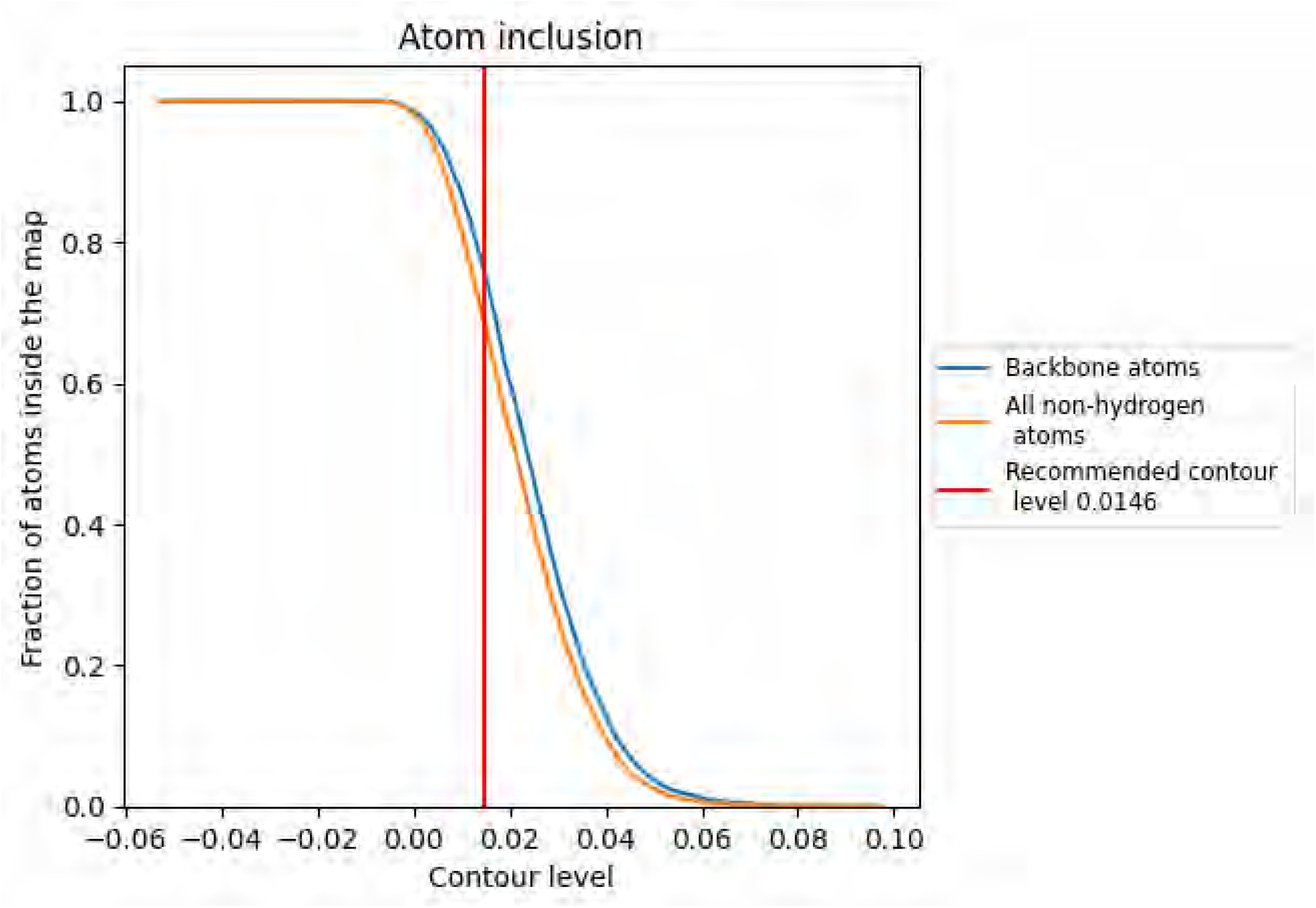

At the recommended contour level, 76% of all backbone atoms, 69% of all non-hydrogen atoms, are inside the map.

